# A single cell atlas of human teeth

**DOI:** 10.1101/2021.02.19.431962

**Authors:** Pierfrancesco Pagella, Laura de Vargas Roditi, Bernd Stadlinger, Andreas E. Moor, Thimios A. Mitsiadis

## Abstract

Teeth exert fundamental functions related to mastication and speech. Despite their great biomedical importance, an overall picture of their cellular and molecular composition is still missing. In this study, we have mapped the transcriptional landscape of the various cell populations that compose human teeth at single-cell resolution, and we analyzed in deeper detail their stem cell populations and their microenvironment. Our study identified great cellular heterogeneity in the dental pulp and the periodontium. Unexpectedly, we found that the molecular signatures of the stem cell populations were very similar, and that their distinctive behavior could be due to substantial differences between their microenvironments. Our findings suggest that the microenvironmental specificity is the potential source for the major functional differences of the stem cells located in the various tooth compartments and open new perspectives towards cell-based dental therapeutic approaches.

## Introduction

Teeth are composed of a unique combination of hard and soft tissues. Enamel, the hardest tissue of the human body, covers the crown of the tooth, and it is supported by a second mineralized tissue, dentin. The central portion of the tooth is occupied by the dental pulp, a highly vascularized and innervated tissue that is lined by odontoblasts, the cells responsible for dentin formation. The tooth is anchored to the surrounding alveolar bone via the periodontium, which absorbs the various shocks associated with mastication and provides tooth stability by continuously remodeling its extracellular matrix, the periodontal ligament (Nanci, 2013). The development of the tooth results from sequential and reciprocal interactions between cells of the oral epithelium and the cranial neural crest-derived mesenchyme (Kollar, 1986; Mitsiadis and Graf, 2009; Nanci, 2013). Oral epithelial cells give rise to ameloblasts that produce enamel. Dental mesenchymal cells give rise to odontoblasts that form the less mineralized tissue of dentin, as well as to the dental pulp (Mitsiadis and Graf, 2009; Nanci, 2013). Dental pulp and periodontal tissues contain mesenchymal stem cells (MSCs), namely the dental pulp stem cells (DPSCs) and periodontal stem cells (PDSCs) (Gronthos et al., 2000; Roguljic et al., 2013). The epithelial cell remnants in the periodontal space upon dental root completion form an additional tooth specific epithelial stem cell population (Athanassiou-Papaefthymiou et al., 2015). DPSCs and PDSCs are multipotent and respond to a plethora of cellular, chemical and physical stimuli to guarantee homeostasis and regeneration of dental tissues. Isolated DPSCs and PDSCs are the subject of intense investigation as possible tools for the regeneration of both dental and non-dental tissues (Chen et al., 2020; Iohara et al., 2011; Lei et al., 2014; Orsini et al., 2018; Ouchi and Nakagawa, 2020; Trubiani et al., 2019; Xuan et al., 2018). In vivo studies aiming at the regeneration of dental pulp and periodontal tissues were however not completely successful (Chen et al., 2020; Xu et al., 2019; Xuan et al., 2018). Indeed, the behavior of these and other stem cell populations is regulated by molecular cues produced in their microenvironment by stromal cells, neurons, vascular-related cells and immune cells, as well as by physical factors such as stiffness, topography and shear stress(Chacon-Martinez et al., 2018; Machado et al., 2016; Oh and Nor, 2015; Oh et al., 2020; Pagella et al., 2015; Rafii et al., 2016; Scadden, 2014; Yang et al., 2017). Much effort has been spent in the last decades to understand the fine composition of tissues, and the cellular and molecular mechanisms that mediate the cross-talk between stem cells and their environment to drive regenerative processes (Blache et al., 2018; Chakrabarti et al., 2018; Lane et al., 2014; Mitsiadis et al., 2017a; Oh et al., 2020; Rafii et al., 2016). Concerning teeth, one recent article reported the single-cell RNA sequencing analysis of mouse dental tissue and the human dental pulp, focusing mostly on the continuously growing mouse incisor and on the conservation between species of cellular populations and features that underlie tooth growth (Krivanek et al., 2020). A second single-cell RNA sequencing analysis study in the continuously erupting mouse incisor identified dental epithelial stem cells subpopulations that are important upon tooth injury and contribute to enamel regeneration (Sharir et al., 2019). Despite the great clinical relevance, the cellular composition of the two main human dental tissues, the dental pulp and the periodontium, has not been investigated in deeper detail.

## Results

We used single cell profiling to elucidate the cellular and molecular composition of human teeth and shed light on fundamental biological questions concerning dental stem cell behavior. We first characterized the cell populations that compose the dental pulp and periodontal tissues in human teeth, and thus we focused on the MSC populations. We isolated pulps from five extracted third molars, dissociated them into single-cell suspensions and proceeded with droplet-based encapsulation (using the 10x Genomics Chromium System) and sequencing. Our analyses yielded a total of 32’378 dental pulp cells (Supplementary Fig. S1). We identified 15 clusters of cells using the graph clustering approach implemented in Seurat v3 (Hafemeister and Satija, 2019) and visualized them using uniform manifold approximation and projection (McInnes et al., 2018) (Figure 1A-D). Our analysis identified a variety of cell populations including mesenchymal stem cells (MSCs), fibroblasts, odontoblasts, endothelial cells (ECs), Schwann cells (ScCs), immune cells, epithelial-like cells and erythrocytes (Figure 1B). MSCs were characterized by the higher expression of *FRZB, NOTCH3, THY1* and *MYH11* (Figure 1C, D, G) (log-fold change of 2.07, 1.24, 1.54 and 1.4 respectively, and adjusted p value <0.001, compared to other cell types in the pulp) and represented on average 12% of the dental pulp tissue (mean proportion = 0.12 and sd = 0.05; Figure 1E). MSCs were localized around the vessels (Figure 1G), where the perivascular niches are formed (Lovschall et al., 2007; Shi and Gronthos, 2003), as well as in the sub-odontoblastic area, which is another potential stem cell niche location in the dental pulp (Mitsiadis and Rahiotis, 2004; Mitsiadis et al., 2003). The fibroblastic compartment composed the bulk of the dental pulp tissue (mean proportion = 0.38 and sd = 0.1; Figure 1E). Different fibroblastic clusters could be identified. Fibroblasts were characterized by the expression of *Collagen*-coding genes (e.g., *COL1A1;* logFC = 0.91 and adjusted p value < 0.001) and *MDK (*logFC = 1.44 and adjusted p value < 0.001), a gene whose expression is restricted to the dental mesenchyme during mouse odontogenesis (Mitsiadis et al., 1995), as well as by the reduced expression of *FRZB* (logFC = −0.76 and adjusted p value < 0.001; Figure 1B-D). One cluster, characterized by the high expression of *Osteomodulin/Osteoadherin* (*OMD*) (Supplementary Fig. S3), represented an intermediate state between MSCs and fibroblasts, with shared gene expression from these two groups. Odontoblasts were characterized by the expression of *Dentin Sialophosphoprotein* (*DSPP*) (Figure 1I) and *Dentin Matrix Acidic Phosphoprotein 1* (*DMP1*), genes encoding for phosphoproteins that constitute essential components of the dentin matrix (D’Souza et al., 1997; Liang et al., 2019). ECs, which constitute important components of the MSCs microenvironment (Rafii et al., 2016), showed a significant degree of heterogeneity (Figure 1B, I; Supplementary Fig. S4). Three well-defined clusters of ECs were detected. A first cluster was characterized by the expression of *EDN1/CLDN5* and represented arterial ECs (Supplementary Fig. S4). A second endothelial cluster was characterized by the expression of *ACKR1/CD234* (Figure 1I; Supplementary Fig. S4) and represented postcapillary and collecting venules. The third main endothelial cluster was characterized by the expression of the *Insulin receptor* (*INSR*) and *RGCC* (Supplementary Fig. S4). Immune cells are part of all healthy tissues and organs (Senovilla et al., 2013) and were also consistently detected in the healthy dental pulp tissues. This cluster mostly consisted of T-cells and macrophages, characterized by the expression of *PTPRC, CD3E,* and *CSF1R* (Figure 1B; Supplementary Fig. S5). Nerve fibers are crucial elements of stem cell niches, as they regulate MSCs functions and fates (Pagella et al., 2015). ScCs formed two clearly distinct clusters of *SOX10*^+^ cells, identified as myelinating *MBP*^+^-ScCs and non-myelinating *GFRA3*^+^-ScCs (Figure 1B, C, J; Supplementary Fig. S6A). *MBP*^+^-ScCs were mostly localized around major nerve fibers entering the dental pulp, while *GFRA3*^+^-ScCs were detected at a distance from nerve fibers and mostly within the sub-odontoblastic regions (Figure 1J), where *NOTCH3*-expressing MSCs were localized. We further identified an epithelial-like cell population within the human dental pulp tissue (Figure 1B, C), in accordance with previous reports in human deciduous teeth (Nam and Lee, 2009). These epithelial cells express Keratin-coding genes such as *KRT14* and *KRT5,* as well as *Stratifin* (*SFN*) (Figure 1C; Supplementary Fig. S6B). We validated the presence of the epithelial cluster within the dental pulp with an immunofluorescent staining against Keratin14 (Figure 1K). We finally identified a population of erythrocytes that is characterized by the presence of the *beta-Hemoglobin*-coding transcript *HBB*.

**Figure 1.**
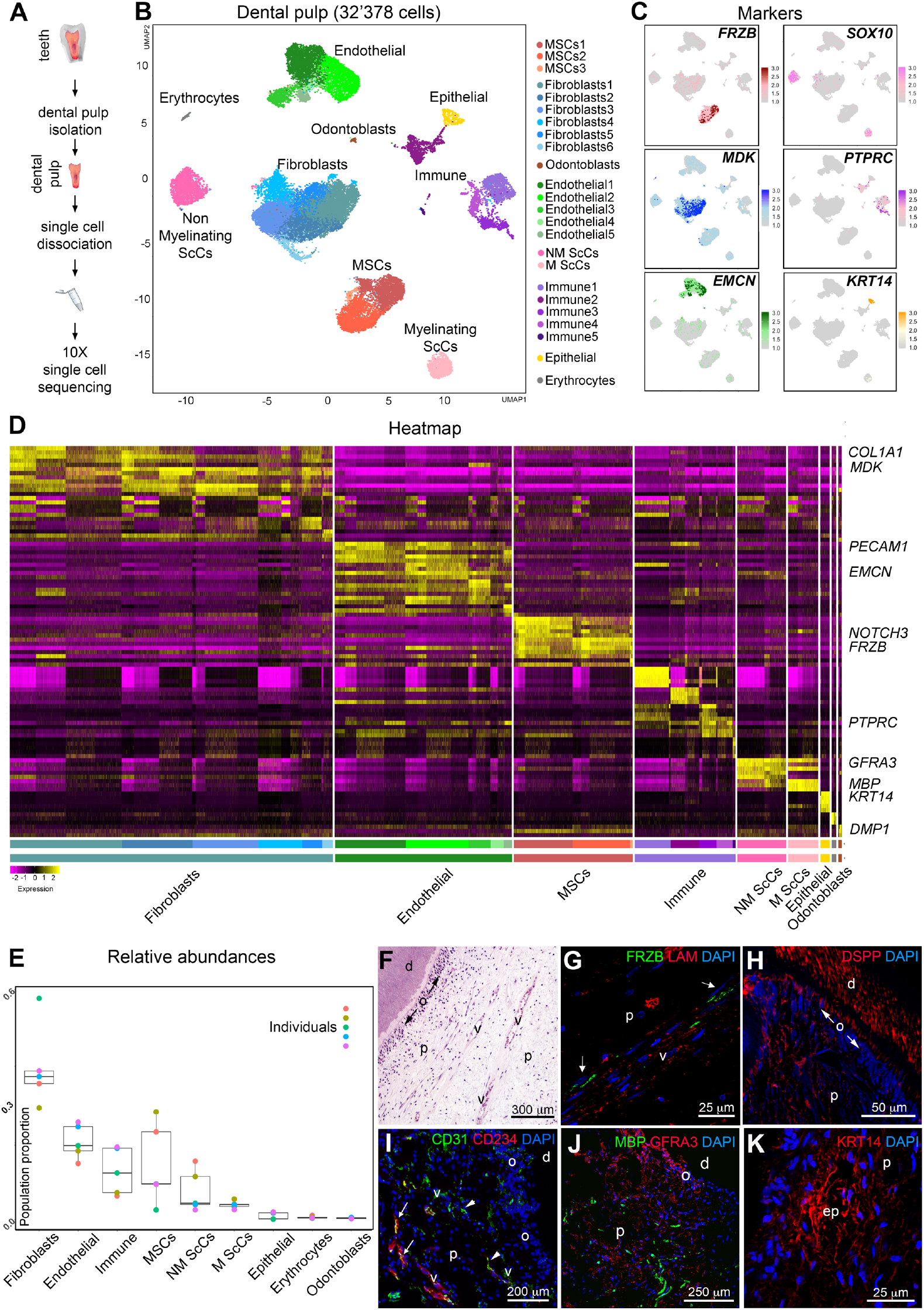
Single cell RNA sequencing analysis of adult healthy human dental pulps. A) Schematic representation of the experimental setup. B) UMAP visualization of color-coded clustering of the dental pulp (n > 32’000 cells). C) Expression of example key genes used for the annotation and the characterization of the clusters. D) Heatmap showing expression of most differentially expressed genes between each cluster and all others. E) Boxplot of relative abundance of main cell types composing the dental pulp from each patient. Boxes illustrate the interquartile range (25th to 75th percentile), the median is shown as the middle band, and the whiskers extend to 1.5 times the interquartile range from the top (or bottom) of the box to the furthest datum within that distance. Any data points beyond that distance are considered outliers. F) Hematoxylin-eosin staining showing the structure of the dental pulp. G) Immunofluorescent staining showing localization of FRZB-expressing (green color, white arrows) MSCs around blood vessels (Laminin-positive, red color). H) Immunofluorescent staining showing localization of DSPP-expressing odontoblasts. I) Immunofluorescent staining showing CD234-expressing (red color) endothelial cells (CD31+, green color). CD31+CD234+ cells are marked by arrows; CD31+CD234-cells are marked by arrowheads. J) Immunofluorescent staining showing localization of MBP-expressing, myelinating Schwann cells (green color) and GFRA3-expressing, non-myelinating Schwann cells (M, red color). K) Immunofluorescent staining showing localization of KRT14-expressing epithelial cells (red color). Blue color: DAPI. Abbreviations: d, dentin; nf, nerve fibers; o, odontoblasts; p, pulp; v, vessels. Scale bars: F, 300 μm; G and K, 25 μm; H, 50 μm; I, 200 μm; J, 250 μm.

We then analyzed the cell populations that compose the human periodontium. We obtained the periodontal tissue by scraping the surface of the apical two-thirds of the roots of five extracted third molars. We dissociated the isolated periodontal tissue to single-cell suspensions and processed them for single cell RNA sequencing (Figure 2A). We obtained a total of 2’883 periodontal cells (Supplementary Fig. S2A) and identified 15 clusters of cells (Figure 2B). MSCs, fibroblasts, ECs, ScCs, immune cells, epithelial-like cells and erythrocytes composed the human periodontal tissue (Figure 2B). Similar to the dental pulp tissue, MSCs represented a large fraction of the periodontium (mean proportion = 0.19, sd = 0.11 and se = 0.05). We detected a cluster of MSCs expressing *FRZB*, *NOTCH3*, *MYH11* and *THY1* (logFC of 1.83, 1.24, 1.47 and 1.61 respectively and adjusted p value < 0.001, compared to other cells in the periodontium; Figure 2B, C). The fibroblastic compartment was defined by cells expressing *MDK (*logFC = 1.25 and adjusted p value <0.001; Figure 2C) and *Collagen*-coding genes such as *COL1A1* (Figure 2B, C; logFC = 3.42 and adjusted p value <0.001). This cluster represented a small fraction of the periodontium (mean proportion = 0.11, sd = 0.08 and se = 0.03; SI Appendix, Fig. S2). ECs were more abundant than fibroblasts and represented a big proportion of the periodontal tissues (mean proportion = 0.19, sd = 0.17 and se = 0.07; Supplementary Fig. S2B). We distinguished two main separate ECs clusters, which were characterized by the expression of *EDN1/CLDN5/CXCL12* and *ACKR1/CD234* (Figure 2B), similar to what observed in the dental pulp (Figure 1B). A cluster of *INSR*/*RGCC*-expressing ECs was observed as an intermediate state between the *EDN1/CLDN5/CXCL1* and *ACKR1/CD234* ECs clusters. ScCs represented a minor population within the periodontium. ScCs expressed *SOX10*, *GFRA3*, *NGF* and *NGFR* (Figure 2B, C; Dataset). The periodontium was characterized by the presence of *PTPRC*^*+*^ immune cells, including T-cells (*CD3E*^*+*^/*CD3D*^*+*^), B-cells (*MZB1*^*+*^), monocytes and macrophages *(CSF1R*^*+*^) (Figure 2C; Dataset). Unexpectedly, we found that the most abundant population of the periodontium consisted of epithelial cells (mean proportion = 0.28, sd = 0.27 and se = 0.12) (Figure 2B-E; Supplementary Fig. S2B). Epithelial cells formed different sub-clusters, characterized by the expression of epithelial genes such as *KRT14, ODAM,* signaling molecules such as *WNT10A*, and specific sets of *Interleukin*-coding genes such as *IL1A* and *IL1B* (Supplementary Fig. S7). Using immunofluorescent staining we showed that the epithelial cells are organized in discrete islets along the entire periodontium (Figure 2H, I). Finally, we identified a small cluster of erythrocytes expressing *HBB* (Figure 2B).

**Figure 2.**
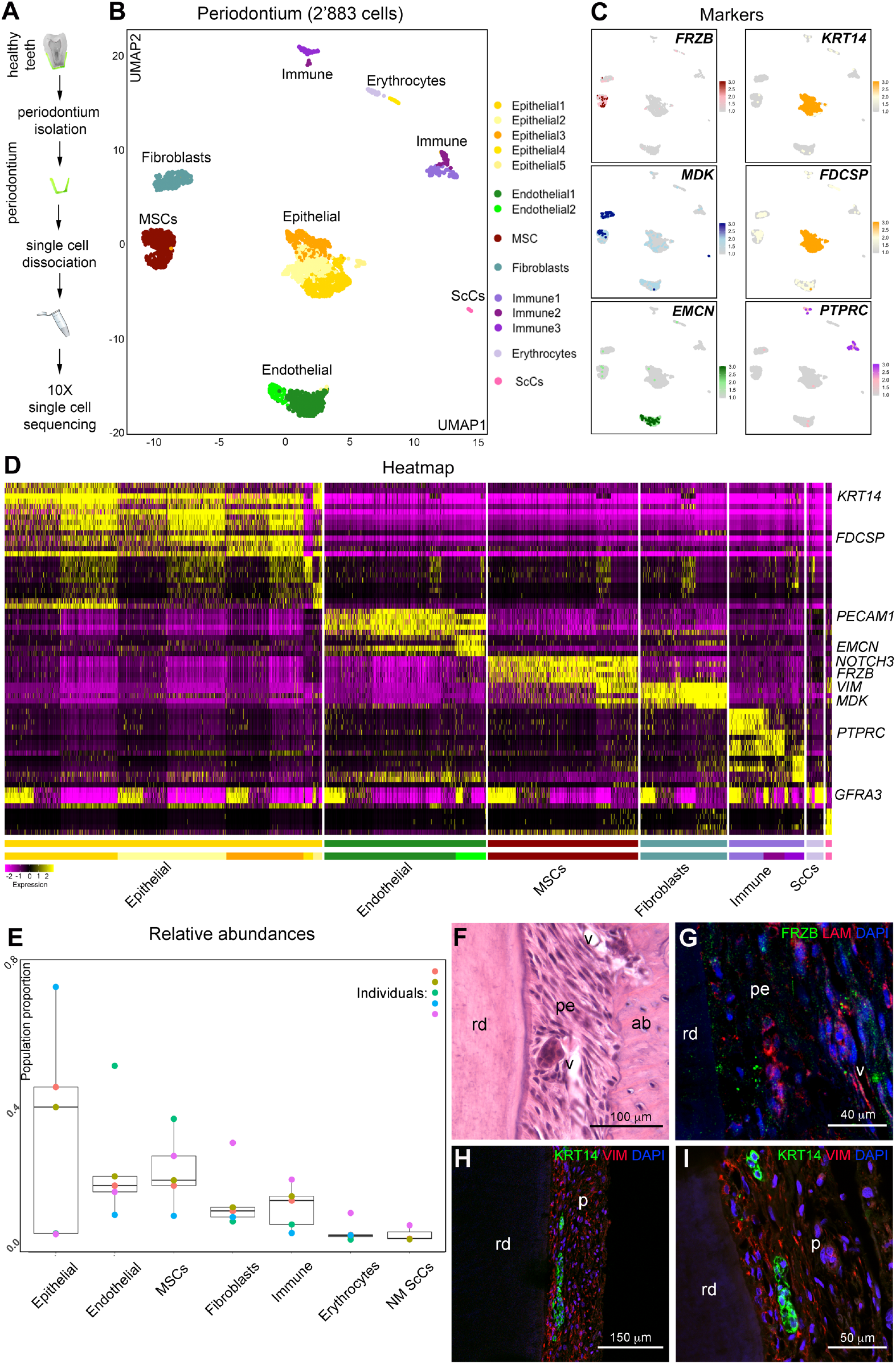
Single cell RNA sequencing analysis of periodontium. A) Schematic representation of the experimental setup. B) UMAP visualization of color-coded clustering of the periodontium (n > 2.800 cells). C) Expression of example key genes used for the annotation and the characterization of the clusters. D) Heatmap showing expression of the gene most differentially expressed between each cluster and all other ones. E) Boxplot of relative abundance of main cell types composing the dental periodontium from each patient. Boxes illustrate the interquartile range (25th to 75th percentile), the median is shown as the middle band, and the whiskers extend to 1.5 times the interquartile range from the top (or bottom) of the box to the furthest datum within that distance. Any data points beyond that distance are considered outliers. F) Hematoxylin-eosin staining showing the structure of the periodontium. G) Immunofluorescent staining showing localization of FRZB-expressing MSCs (green color). Red color: laminin, marking blood vessels; blue color: DAPI. H) Immunofluorescent staining showing localization of KRT14-expressing epithelial cells (green color) and Vimentin-expressing (VIM) mesenchymal cells (red color) along the periodontium. Blue color: DAPI. I) Higher magnification of H. Abbreviations: ab, alveolar bone; lam, laminin; pe, periodontium; rd, root dentin; v, vessel. Scale bars: F, 100 μm; G, 40 μm; I, 50 μm; H, 150 μm.

The establishment of this map allows for further analyses and comparisons at the molecular level between the dental pulp and periodontium (Supplementary Fig. S8A, B). We thus first focused on the molecular analysis of the MSCs clusters detected in the dental pulp and periodontal tissues. In both tissues, MSCs were characterized by the expression of *FRZB* and *NOTCH3* (logFC = 2.05 and 1.27 and p values <0.001; Figure 3A, C; Supplementary Fig. S8C, D). We then analyzed the composition of the dental pulp and periodontal MSCs clusters in deeper detail. Upon separate sub-clustering of the *NOTCH3*^*+*^*FRZB*^*+*^ pulp and periodontal MSCs we identified three major MSCs subpopulations (Figure 3A-C; Supplementary Fig. S8C, D). Unexpectedly, the main dental pulp and periodontal MSCs populations exhibited very similar molecular signatures. Both compartments contained two main MSCs clusters characterized by increased expression of *MYH11* (logFC = 2.00 and p value < 0.001) and *THY1* (logFC = 1.63 and p value < 0.001), respectively, when compared to all other clusters (Figure 3B, D; Supplementary Fig. S8C). We detected a second *THY1*-positive (and *MYH11*-negative) MSC cluster, with increased expression of *CCL2* (logFC = 3.46 and p value < 0.001 when compared to other clusters; Figure 3B, D). The *CCL2*^*+*^ MSCs cluster also expressed genes associated with the remodeling of the extracellular matrix, such as *TNC* (Tenascin C) (Supplementary Fig. S8D).

**Figure 3.**
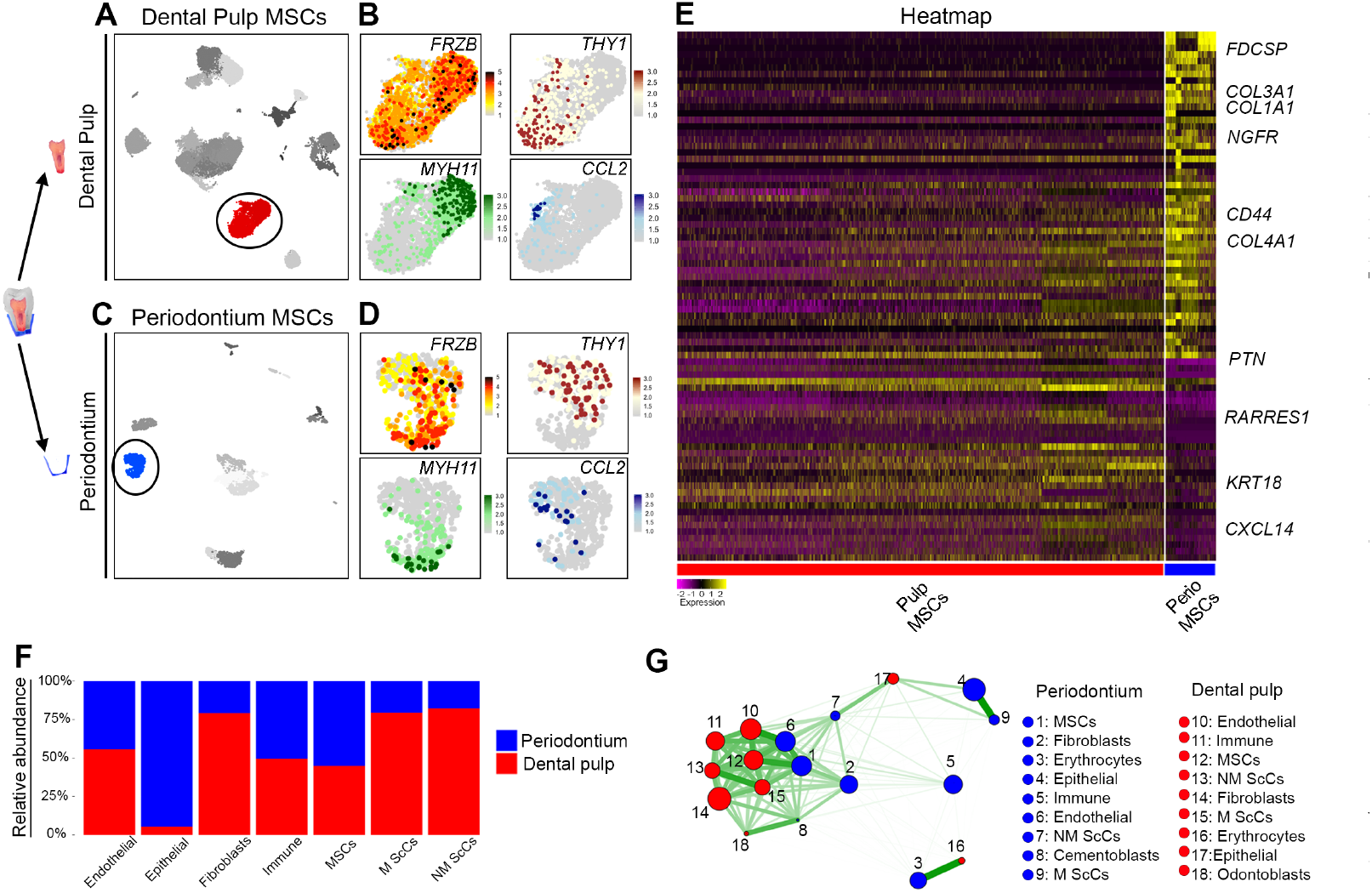
Comparative analysis of MSCs compartment and their microenvironment in the pulp and the periodontium. A) UMAP visualization of pulp clusters, highlighting the MSCs compartment. B) Feature plots showing genes that characterize the main MSCs subclusters within the pulp. C) UMAP visualization of periodontium clusters, highlighting the MSCs compartment. D) Feature plots showing genes that characterize the main MSCs subclusters within the periodontium. E) Heatmap showing differential gene expression between periodontium and pulp MSCs. See SI Appendix, Table S1, for logFC and adjusted p value. F) Comparison of the relative abundance of the different cell types composing the pulp and the periodontium. Epithelial cells are the most abundant cell type in the periodontium, while fibroblasts constitute the most abundant cluster in the dental pulp. G) Extended Jaccard similarity plot between the various periodontium and pulp clusters using the top three thousand differentially expressed genes. See SI Appendix, Table S2, for extended Jaccard similarity ranking.

Next, we merged the dental pulp and periodontium datasets and jointly clustered them to compare the transcriptomes of their MSCs (Supplementary Fig. S8). We detected gene expression log-fold changes higher than 0.25 in only 333 genes and as few as 33 genes with a logFC higher than 1(p values <0.05, Figure 3E; Supplementary Fig. S8B; Table S1). MSCs from the two tissues showed no significant differences in the expression of the already mentioned *NOTCH3*, *FRZB*, *THY1*, *MYH11* (Figure 1, 2) as well the other stem cell markers *MCAM/CD146*, *RGS5, ACTA2* and *ID4* (Supplementary Fig. S8D). Some genes were significantly more expressed in periodontal MSCs than in the pulp, such as *CCL2* (logFC = 0.78 and p< 0.001), and those coding for Collagens (e.g., *COL3A1*, *COL1A1*, *COL6A1*, *COL6A3*, *COL4A1*) (logFC = 1.65, 1.59, 1.02, 0.70, 0.86, respectively and adjusted p values <0.001; Figure 3E; Table S1, S3; Supplementary Fig. S9). Periodontal MSCs were also characterized by higher expression of *SPARC*/*Osteonectin,* a secreted molecule fundamental for the regulation of periodontal homeostasis and collagen content (logFC = 1.00 and p value <0.001; Figure 3E; Table S1, S6). In contrast to the periodontal MSCs, dental pulp MSCs expressed higher levels of *CXCL14* and *RARRES1* (logFC = 2.04 and 1.00, respectively and p values <0.001; Figure 3E; Table S1). Surprisingly, dental pulp MSCs strongly expressed *KRT18,* a gene previously reported to be exclusively expressed in cells of single-layered and pseudo-stratified epithelia (logFC = 1.46, p value <0.001; Table S1, Supplementary Fig. S11). We further estimated the pairwise extended jaccard similarity for all cell types present in the periodontium and dental pulp and ranked these pairwise similarities. This analysis revealed that the three most similar cell types between the periodontium and the dental pulp were, in order, Endothelial cells, Erythrocytes and MSCs (Figure 3G and Table S2).

We then compared the two specific MSC niches in the dental pulp and the periodontium (Figure 3F; Supplementary Fig. S2, S8). We observed that their cell compositions diverged in relative proportion for certain cell types, mainly the fibroblastic and epithelial compartments. Fibroblasts represented the most abundant cell population within the dental pulp, while in the periodontium the proportion of fibroblasts was considerably lower (mean dental pulp = 0.38 and se = 0.04; mean periodontium: 0.11 and se = 0.03. Figure 3F; Supplementary Fig. S2). Likely due to the high variability of scRNA-seq, it is not possible to statistically confirm this difference using our dataset. Genes coding for Collagens and Matrix Metalloproteases (MMPs) were highly expressed by periodontal fibroblasts and MSCs (Supplementary Fig. S9, S10 and Table S3, S5) when compared to their pulp counterparts. Interestingly, genes coding for bone-specific proteins, such as Osteonectin (*SPARC*), Osteocalcin (*BGLAP*) and Bone Sialophosphoprotein (*BSP*), were expressed by the periodontal fibroblasts (Supplementary Fig. S12 and Table S6). Periodontal fibroblasts also expressed MGP (Matrix Gla Protein), a potent inhibitor of mineralization (Supplementary Fig. S12, Table S6). The periodontium was characterized by a larger proportion of cells expressing epithelial cell markers such as *KRT5* and *KRT14* (Supplementary Fig. S2B). As in the case of fibroblasts, it was not possible to statistically confirm this difference in proportion. These periodontal epithelial-like cells expressed different sets of *Keratin*-coding genes when compared to those of the dental pulp (Supplementary Fig. S11, Table S4). In the periodontium, Keratin-coding genes such as *Krt14*, *Krt17*, and *Krt19* were not exclusively expressed by epithelial cells, but also significantly enriched in fibroblasts and ScCs (Supplementary Fig. S11 and Table S4). The periodontal epithelial-like cells also expressed genes encoding for signaling molecules such as FDCSP (Follicular Dendritic Cells Secreted Protein) and WNT10A (Figure 2D; Supplementary Fig. S7). We also found that in the periodontium the MSCs expressed significantly higher levels of *Collagen*-coding genes (e.g., *COL1A1*, *COL3A1*, *COL6A1;* Supplementary Fig. S9A).

We analyzed the overall dynamics and differentiation trajectories of dental pulp and periodontal MSCs by Velocity (Supplementary Fig. S13). We did not identify major differentiation trajectories between different cell types neither in the dental pulp nor in the periodontium. In the dental pulp, endothelial cells showed the most dynamic behavior, while only minor differentiation trajectories were identified within most dental pulp cell populations (Supplementary Fig. S13). In the periodontium, epithelial-like cells, fibroblasts and MSCs displayed dynamic behaviors (Supplementary Fig. S13). Periodontal MSCs showed a directional gene expression trajectory from the *MYH11*^+^ to the *THY1*^*+*^ sub-cluster (Supplementary Fig. S13). Periodontal *THY1*^*+*^ MSCs co-expressed genes that characterize the fibroblastic compartment, such as the *Collagen*-coding genes, *MMP14* and *SPARC* (Supplementary Fig. S9, S12). *MYH11*^+^ cells might thus constitute the most undifferentiated pool of MSCs within the periodontal tissue, while the *THY1*^*+*^ sub-cluster could represent MSCs directed towards the fibroblastic fate.

## Discussion

Understanding the fine composition of human organs is of paramount importance to develop regenerative therapies. In particular, unraveling the composition of stem cell populations and their niches is fundamental to drive regenerative processes towards the reconstitution of fully functional tissues and organs. This study revealed that MSCs in the human dental pulp and periodontium are characterized by the expression of *FRZB* and *NOTCH3, THY1* and *MYH11. Frzb* had already been shown to mark periodontal ligament cells from very early developmental stages (Mitsiadis et al., 2017b), while its expression in the dental pulp had not yet been reported. Previous studies have also shown that *Notch3* is expressed in perivascular MSCs both in dental and non-dental tissues (Jamal et al., 2015; Lovschall et al., 2007; Wang et al., 2014). Both dental pulp and periodontium MSCs can be subdivided into subpopulations characterized by the expression of the same specific markers *MYH11*, *THY1*, and *CCL2*. *THY1*/ *CD90* is a general marker of human mesenchymal stem cells, and it is vastly used to sort human dental pulp stem cells (Dominici et al., 2006; Ledesma-Martinez et al., 2016; Sharpe, 2016). *MYH11* is mostly known to be expressed in smooth muscle cells, and in our datasets, it was generally co-expressed with *ACTA2* (α-smooth muscle actin), recently found to play an important role in MSCs cell fate specification (Talele et al., 2015). *CCL2* codes for the chemokine ligand 2, and its expression in MSCs was shown to be a key mediator of their immunomodulatory properties (Giri et al., 2020). Expression of these markers in the dental pulp MSCs is in accordance with the datasets reported in a recent work (Krivanek et al., 2020), while the existence of three distinct MSC subclusters, both in the dental pulp and periodontium, was not reported before. Beyond the expression of these markers, dental MSCs show an overall striking homogeneity, in contrast to current assumptions (Hakki et al., 2015; Lei et al., 2014; Otabe et al., 2012). Indeed, previous studies have shown that although dental pulp and periodontal MSCs possess similar differentiation potentials in generating adipoblasts, myoblasts, chondroblasts, and neurons, their efficacies in forming bone tissues differ (Bai et al., 2010; d’Aquino et al., 2011; Schiraldi et al., 2012; Yagyuu et al., 2010). Human dental pulp and periodontal MSCs do not differ in their specific migratory behavior when cultured separately *in vitro* (Schiraldi et al., 2012). However, when these two cell types are co-cultured, the periodontal MSCs quickly spread and directionally migrate towards the dental pulp MSCs, which exhibit limited proliferative and migratory capabilities (Schiraldi et al., 2012). MSCs proliferation and directional migration cues are generally produced by the target tissue, as well as by direct contacts established through the interactions of MSCs with cells composing their niches (Schiraldi et al., 2012; Shellard and Mayor, 2019). The divergent behavior of these MSCs, both in migration and in differentiation, could be due to their interaction with different environments rather than due to intrinsic differences. Our results support that dental MSCs homogeneity is counteracted by a great divergence in their niches. In our samples, the dental pulp was composed mostly by fibroblasts, while epithelial cells constituted the most abundant cluster in the periodontium. Fibroblasts and epithelial cells within the dental pulp and the periodontium also expressed very different sets of molecules that could modulate MSCs behavior. Genes coding for Collagens and Matrix Metalloproteases (MMPs), as well as genes encoding for regulators of mineralization such as Osteonectin, were highly expressed by periodontal fibroblasts and MSCs when compared to their pulp equivalent. Osteonectin is known to regulate Ca^2+^ deposition during bone formation, but in the periodontium its function is essential for proper collagen turnover and organization (Luan et al., 2007). Periodontal fibroblasts also expressed MGP (Matrix Gla Protein), a potent inhibitor of mineralization (Kaipatur et al., 2008). The most abundant periodontal cell type is represented by epithelial-like cells. These periodontal epithelial-like cells expressed genes encoding for signaling molecules such as FDCSP (Follicular Dendritic Cells Secreted Protein) and WNT10A, which exert fundamental roles in the modulation of periodontal MSCs proliferation and differentiation (Wei et al., 2011; Xu et al., 2017; Yu et al., 2020). Epithelial cells from the periodontium have been long proposed to constitute a dental epithelial stem cells population, with potential to generate tooth-associated hard tissues such as enamel, dentin and alveolar bone (Athanassiou-Papaefthymiou et al., 2015; Tsunematsu et al., 2016). We showed that these cells also have signaling properties that could influence the behavior of periodontal MSCs. Overall, the cellular and molecular signature of the periodontium identified in this study was indicative of its continuous and dynamic remodeling, which is tightly linked to the masticatory function of the teeth, and that requires continuous collagen secretion, extracellular matrix remodeling, and inhibition of mineralization (Takimoto et al., 2015). Taken together, these great cellular and molecular differences in the microenvironment of the dental pulp and periodontium constitute strong tissue-specific traits that can be indicative of a microenvironment that privileges MSCs differentiation towards a fibroblastic-like fate in the periodontium, as opposed to the dental pulp microenvironment, which favors the osteogenic fate of MSCs. Two recent articles described the single-cell RNA sequencing analysis of dental tissues (Krivanek et al., 2020; Sharir et al., 2019). One study identified the main cell types that compose the dental pulp and compared their behavior in mice and humans, and between human adult and erupting teeth (Krivanek et al., 2020). This work showed that basic features underlying tooth growth, such as lineage hierarchy between *Smoc2*^*−*^ and *Smoc2*^*+*^ cells, are conserved between mice and humans (Krivanek et al., 2020). The datasets concerning the human dental pulp presented in this work are in general agreement with our data. Our results provide a significantly more resolved analysis, in which we identified not only the major cell types present within the dental pulp and the periodontium, but also their heterogeneity. In a second study, the authors performed single cell RNA sequencing analysis of the epithelium of the continuously growing mouse incisor and revealed the role of Notch1-expressing stem cells showing that these cells are responsive to tooth injury and contribute to enamel regeneration (Sharir et al., 2019). Overall, these studies are complementary to our work, as they focused mostly on mouse teeth, while they did not investigate in detail the cell types that compose the human dental pulp and the human periodontium.

Taken together, our findings provide a thorough investigation of the human pulp and periodontal tissues at single-cell resolution, thus representing the basis for future research involving cell-based regenerative treatments.

## Acknowledgements

We thank Dr. Emilio Yangüez and the Functional Genomics Center Zurich (ETH / University of Zurich) for the processing of the samples for single cell RNA sequencing. We thank Ms. Kendra Wernlé and Ms. Madeline Fellner (Institute for Oral Biology, University of Zurich) for technical assistance. Imaging was performed with equipment maintained by the Center for Microscopy and Image Analysis, University of Zurich. This work was financially supported by the University of Zurich and by the Swiss National Science Foundation (310030_197782).

## Author contributions

Conceptualization: T.M., A.M., P.P. Methodology: T.M, A.M., P.P., L.V.R., B.S. Data analysis: A.M., L.V.R. Validation: T.M., A.M., L.V.R., P.P. Formal analysis: P.P., L.V.R., A.M., T.M. Investigation: P.P., L.V.R. Resources: T.M., A.M. Data curation: L.V.R., A.M. Writing – original draft: P.P., T.M. Writing – Review & Editing: P.P., L.V.R., B.S., A.M., T.M. Visualization: P.P., L.V.R., A.M., T.M. Supervision: T.M., A.M. Project administration: T.M., A.M. Funding acquisition: T.M.

## Materials and Methods

### Data and code availability

Data can be found at GEO; Access number: GSE161267. All code is publicly available at: https://github.com/TheMoorLab/Tooth

#### Isolation of cells from the dental pulp and the periodontium for single cell RNA sequencing

The procedure for the collection of anonymized human dental pulp and periodontal cells at the Center of Dental Medicine (ZZM) of the University of Zurich was approved by the Ethic Commission of the Kanton of Zurich (reference number 2012-0588) and the patients gave their written informed consent. Samples were obtained in fully anonymized form from patients of 18-35 years of age. Tooth extractions were performed by professional dentists at the Oral Surgery department of ZZM of the University of Zurich. Evaluation of the health status of the tooth was done post-extraction, upon direct observation of the specimen. All procedures were performed in accordance with the current guidelines. Teeth were collected immediately after extraction and preserved in sterile NaCl 0.9%, on ice for the time needed to transfer them from the clinic to the processing laboratory (< 10 minutes). The periodontium was isolated by scratching the lower two-thirds of the root of the teeth with a surgical scalpel directly into a Petri dish filled with sterile, cold Hank’s Balanced Salt Solution (HBSS; Thermo Fisher Scientific, Reinach, Switzerland). The upper-third of the root was excluded to minimize contamination from the gingival epithelium. The cleansed tooth was then carefully wiped with 70% ethanol. The tooth was then cracked with a press, and carefully opened with forceps. The dental pulp was then removed from the tooth with a separate set of instruments, placed in a Petri dish filled with cold HBSS and minced into small pieces (< 2 mm diameter). Thereafter, periodontal and pulp tissues were transferred in falcon tubes filled with HBSS, centrifuged at 4°C, 300g, for 10 minutes. Tissues were digested in 10 mL Collagenase P 5 U/mL (11 213 873 001, Sigma Aldrich, Buchs, Switzerland) for 40 minutes at 37°C. After digestion, samples were disaggregated by pipetting, filtered through a 70 μm cell strainer, and resuspended in HBSS + 0.002% Bovine Serum Albumin (BSA; 0163.2, Roth AG, Arlesheim, Switzerland).

#### Single-cell RNA sequencing (scRNA-seq) using 10X Genomics Platform

The quality and concentration of the single cell preparations were evaluated using a hemocytometer in a Leica DM IL LED microscope and adjusted to 1’000 cells/μl. 10’000 cells per sample were loaded into the 10X Chromium controller and library preparation was performed according to the manufacturer’s indications (single cell 3’ v2 or v3 protocol). The resulting libraries were sequenced in an Illumina NovaSeq sequencer according to 10X Genomics recommendations (paired end reads, R1=26, i7=8, R2=98) to a depth of around 50.000 reads per cell.

#### Computational analysis

Velocity analysis was performed using scVelo (Bergen et al., 2019) and Python v3.6. Velocity was only calculated for patients’ samples sequenced with 10X v3. All other data analysis was performed using Seurat v3 (Stuart et al., 2019) and R version 3.6.4. Clusters were visualized using uniform manifold approximation and projection (UMAP) (McInnes et al., 2018). Dental pulp and periodontium data were initially analyzed separately. Data was scaled and transformed using SCTransform (Hafemeister and Satija, 2019) for variance stabilization. Analysis of merged dental pulp and periodontium data was performed by integrating data with R package Harmony (Korsunsky et al., 2019) to cluster data into cell types. Any subsequent analysis was done using raw data and not data transformed after integration. In particular, all statistical analysis of differential expression was performed on unintegrated and untransformed data as both could lead to dependencies in the data rendering the assumption of independence of the statistical test void. Differential expression analysis was performed using the Wilcoxon rank sum test. All p values reported were adjusted for multiple comparisons using the Bonferroni correction. The extended Jaccard similarity was computed on the top three thousand differentially expressed genes across the two datasets (pulp and perio samples).

#### Processing of human teeth for immunofluorescent staining

Teeth used for histological analysis and immunostaining were immediately fixed by immersion in paraformaldehyde 4% (PFA 4%) for 24 hours, then decalcified in Morse’s solution for 8 weeks, dehydrated, embedded in paraffin, and serially sectioned at 5 μm. From a subset of teeth, the dental pulp was immediately extracted and fixed in PFA 4% for 2 hours. The specimens were then incubated in Sucrose 30%, embedded in Tissue Tek® O.C.T.TM (4583, Sakura, Alphen aan den Rijn, Netherlands), and serially sectioned at 10 μm.

#### Immunostaining

Paraffin sections were rehydrated by incubation in Xylol followed by a series of Ethanol solutions (100% to 30%) and distilled H_2_O. Cryosections were let dry at room temperature for 1 hour and then washed with PBS before immunostaining. Cells used for immunofluorescent staining were first fixed in PFA 4% for 15 minutes at 4°C. Thereafter, specimens were blocked with PBS supplemented with 2% Fetal Bovine Serum (FBS) and incubated with primary antibodies for 1 hour at room temperature. The following primary antibodies were used: Rabbit anti-Thy1/CD90 (1:100; 555595, BD, Eysin, Switzerland), Rabbit anti-Keratin 14 (1:500; PRB-155P, BioLegend, San Diego, CA, U.S.A.), Mouse anti-Vimentin (1:100; M0725, Dako, Baar, Switzerland), Mouse anti-FRZB (1:50, LS-B6898-50, LSBio, Seattle, WA, U.S.A.), Rabbit anti-Dentin Sialophosphoprotein (DSPP) (1:100, ENH083, Kerafast, Boston, MA, U.S.A.), Rabbit anti-Laminin (1:20; ab11575, Abcam, Cambridge, United Kingdom), rabbit anti-Neurofilament (1:100; 2837S, Cell Signaling Technology, Danvers, MA, U.S.A.), anti-MBP (1:200; MAB386, Millipore), anti-CD31 (1:50; ab28364, Abcam, Cambridge, UK), anti-FABP4 (1:50; ab92501, Abcam, Cambridge, UK), anti-Endothelin 1 (1:100; MA3005, Thermo Fisher Scientific, Reinach, Switzerland), anti-CD234 (1:50; 566424, BD, Eysin Switzerland). The sections were then incubated with Fluorochrome-conjugated secondary antibodies for 1 hour at room temperature at dark. The following secondary antibodies were used: Alexa-568 Donkey anti-Rabbit (1:500; A10042, Thermo Fisher Scientific, Reinach, Switzerland), Alexa-488 Chicken anti-Goat (1:500; A-21467, Thermo Fisher Scientific, Reinach, Switzerland), Alexa-488 Goat anti-Rabbit (1:500; A32731, Thermo Fisher Scientific, Reinach, Switzerland), Alexa-568 Goat anti-Rat (1:500; A-11077, Thermo Fisher Scientific, Reinach, Switzerland). DAPI (4’,6-Diamidino-2-Phenylindole; D1306, Thermo Fisher Scientific, Reinach, Switzerland) was then used for nuclear staining. After immunofluorescent staining, samples were mounted in ProLong™ Diamond Antifade Mountant (P36965, Thermo Fisher Scientific, Reinach, Switzerland), and imaged with a Leica SP8 Inverted Confocal Laser Scanning Microscope (Leica Microsystems-Schweiz AG, Heerbrugg, Switzerland).

