## Supplementary Figures for "A single cell atlas of human teeth"

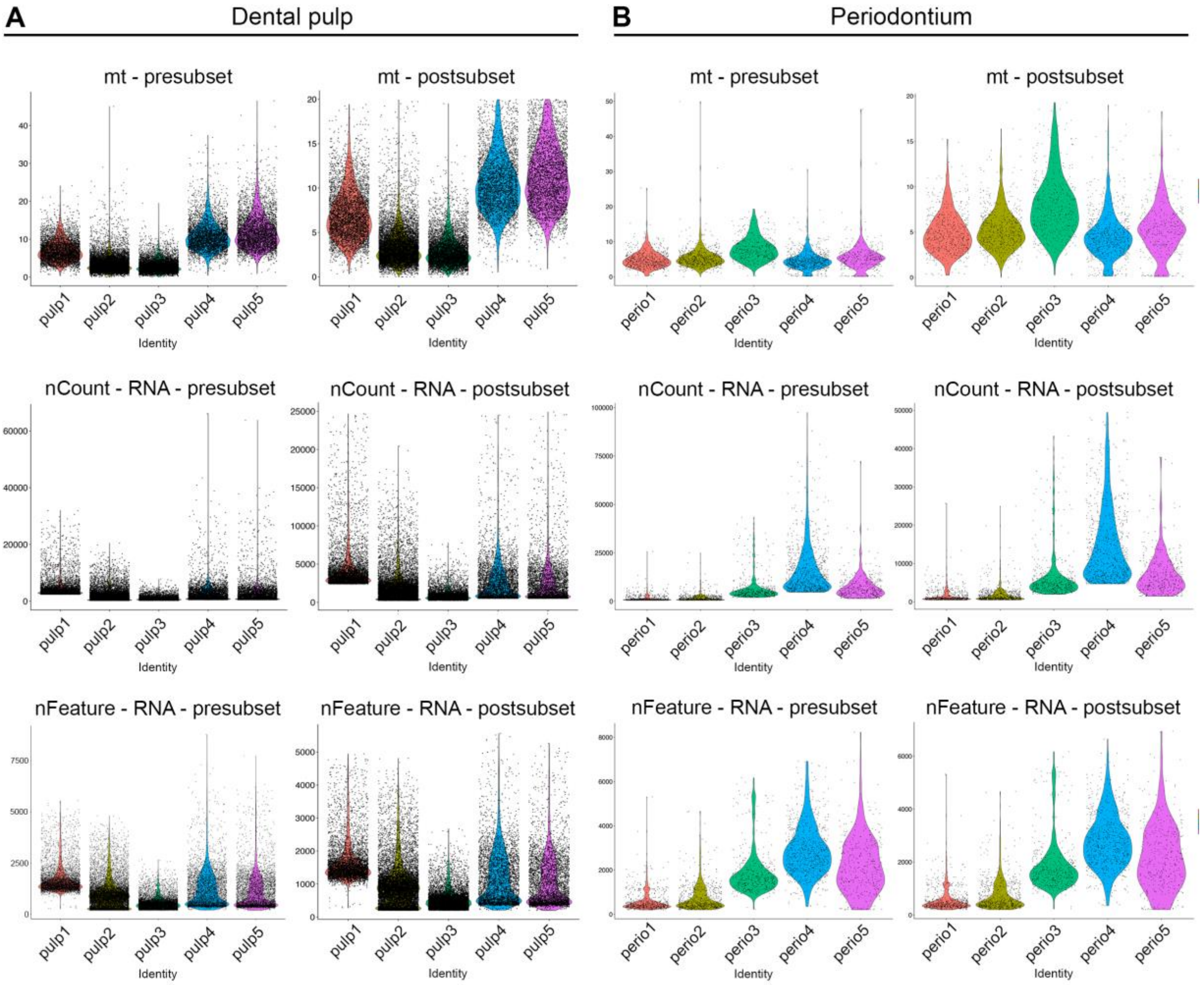


**Fig. S1. Quality control and pre-processing of single-cell pulp (A) and periodontium (B) data.** Violin plots illustrate distribution of percentage of mitochondrial genes (mt), number of UMI counts (nCount) and number of genes with at least one UMI count (nFeature) per cell prior and after subsetting cells according to the following quality control measures: cells with a percentage of mitochondrial genes above 20 were excluded, as well as cells with less than 200 genes. Healthy pulp and periodontal cells with UMI counts above 25'000 and 50'000, respectively were also excluded.

**A** Samples details

| | Dental pulp | Periodontium |
|---|---|---|
| **number of samples** | 5 | 5 |
| **total number of cells** | 32378 | 2883 |
| **median number of cells per sample** | 5100 | 615 |
| **mean number of cells per sample** | 6475.6 | 576.6 |
| **sd number of cells** | 3022.7 | 152.9 |
| **median genes** | 830 | 1333 |
| **mean genes** | 1017.9 | 1612.5 |
| **sd genes** | 732.7 | 1255.8 |

**B** Relative abundance by cell type

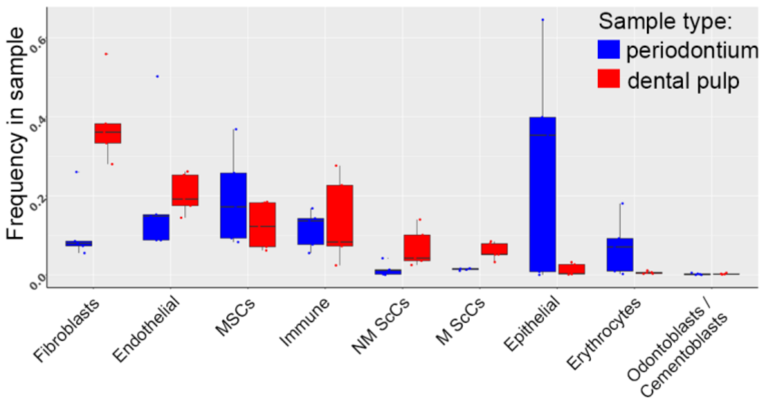


**Fig. S2. Quantitative details of dental pulp and periodontal samples.** A) Sample size and statistics. B) Relative abundance of cell types in the pulp (red) and periodontium (blue); Boxes illustrate the interquartile range (25th to 75th percentile), the median is shown as the middle band, and the whiskers extend to 1.5 times the interquartile range from the top (or bottom) of the box to the furthest datum within that distance. Any data points beyond that distance are considered outliers.

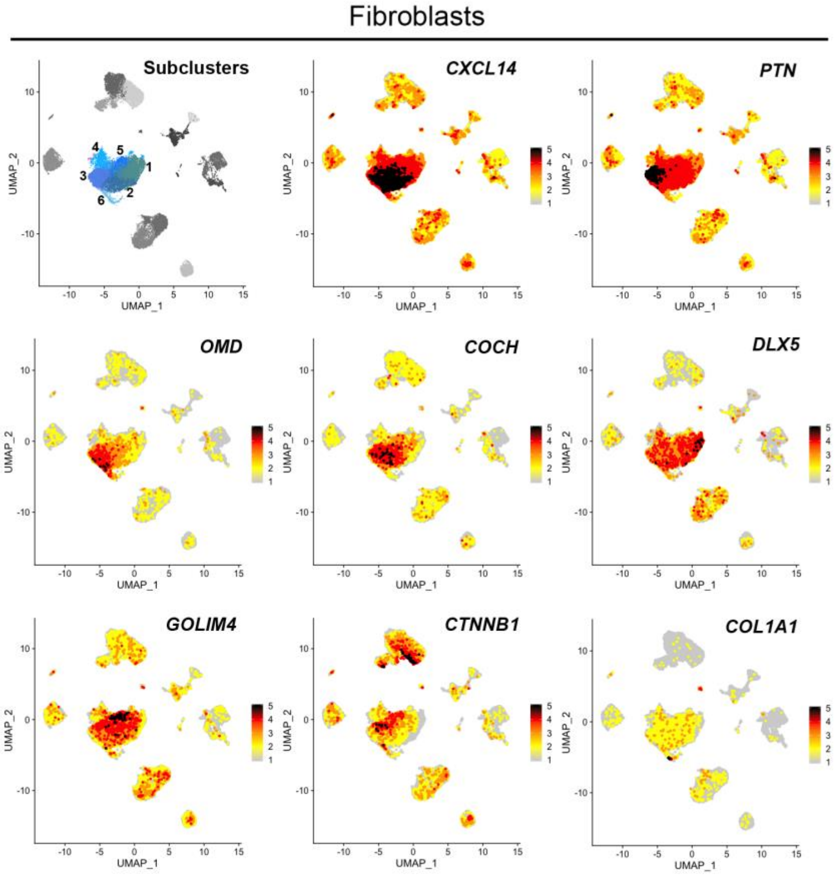


**Fig. S3. Feature plots showing the expression of genes characterizing specific subclusters of dental pulp fibroblasts.** 5 subclusters could be identified within fibroblasts. CXCL14 was generally expressed by fibroblasts, and it was particularly enriched in subclusters 2 and 3. CXCL14 expression is associated with angiogenic potential and overall chemoattractant properties (Hayashi et al., 2015). Subcluster 3 showed higher expression of the dental mesenchyme marker PTN (pleiotrophin), which is associated with odontoblastic differentiation potential (Mitsiadis et al., 1995). Cluster 6 was characterized by higher expression of Osteomodulin/Osteoadherin (OMD), a modulator of mineralization (Buchaille et al., 2000; Lin et al., 2019). This same cluster also showed particularly high expression of COL1A1. Clusters 2 and 6 expressed high levels of COCH, which encodes for a protein involved in mechano-sensation (Goel et al., 2012). Cluster 1 expressed high levels of DLX5, while cluster 5 was characterized by high expression of CTNNB1 and GOLIM4. CTNNB1 codes for b-catenin, key mediator of WNT signaling (Mosimann et al., 2009).

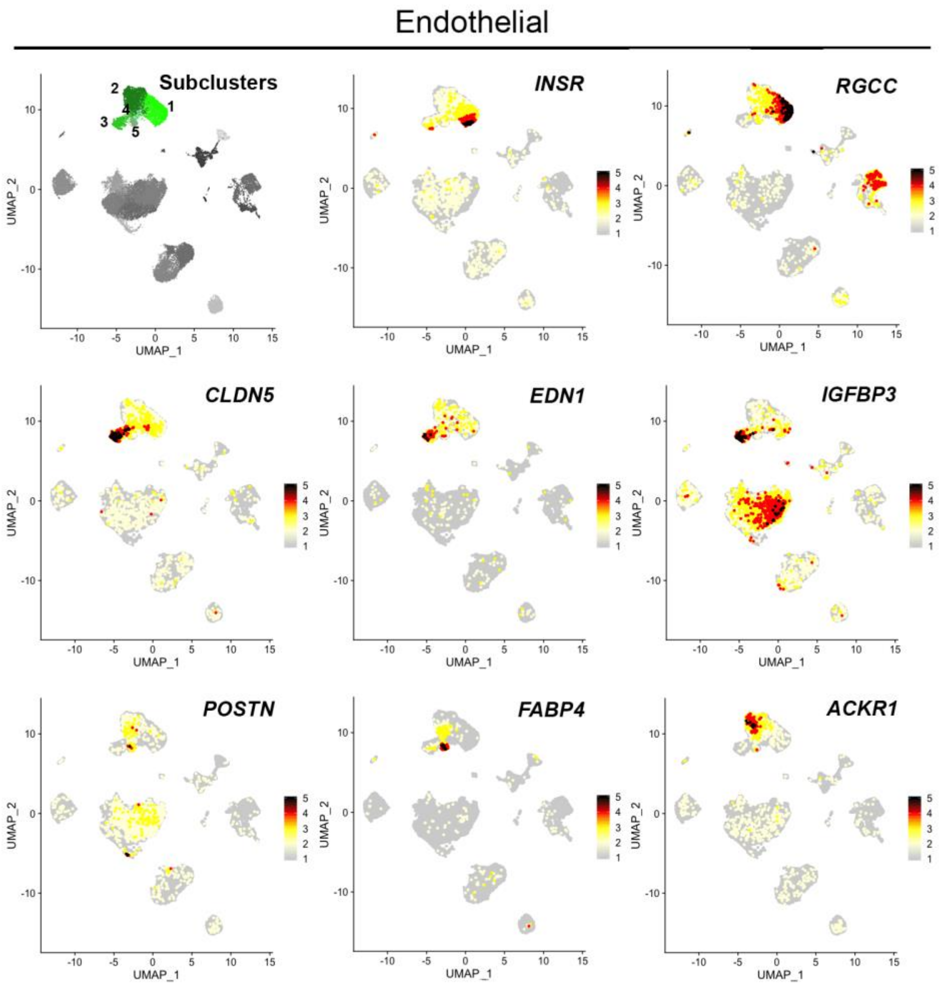


**Fig. S4. Feature plots showing the expression of genes characterizing specific subclusters of dental pulp endothelial cells (ECs).** INSR (Insulin Receptor) and RGCC marked ECs from subcluster 1. Expression of INSR suggests a role for these cells in modulating Glucose and Insulin metabolism within the dental pulp, while RGCC expression is usually observed in actively cycling cells (Konishi et al., 2017; Kubota et al., 2011). CLDN5 (claudin 5), EDN1 (endothelin 1) and IGFBP3 were enriched in ECs from cluster 3. These cells co-expressed the arterial markers GJA5 and EFNB2 (Mukouyama et al., 2002; Shin et al., 2001), thus indicating that this cluster represents arterial ECs. Cluster 5 was characterized by the high expression of POSTN (Periostin) and FABP4, both associated to angiogenic and pro-survival processes in endothelial cells (Elmasri et al., 2012; Hu et al., 2016). Cluster 2 was marked by the expression of ACKR1/CD234, which has been proposed as a marker for postcapillary and collecting venules in mice (Thiriot et al., 2017).

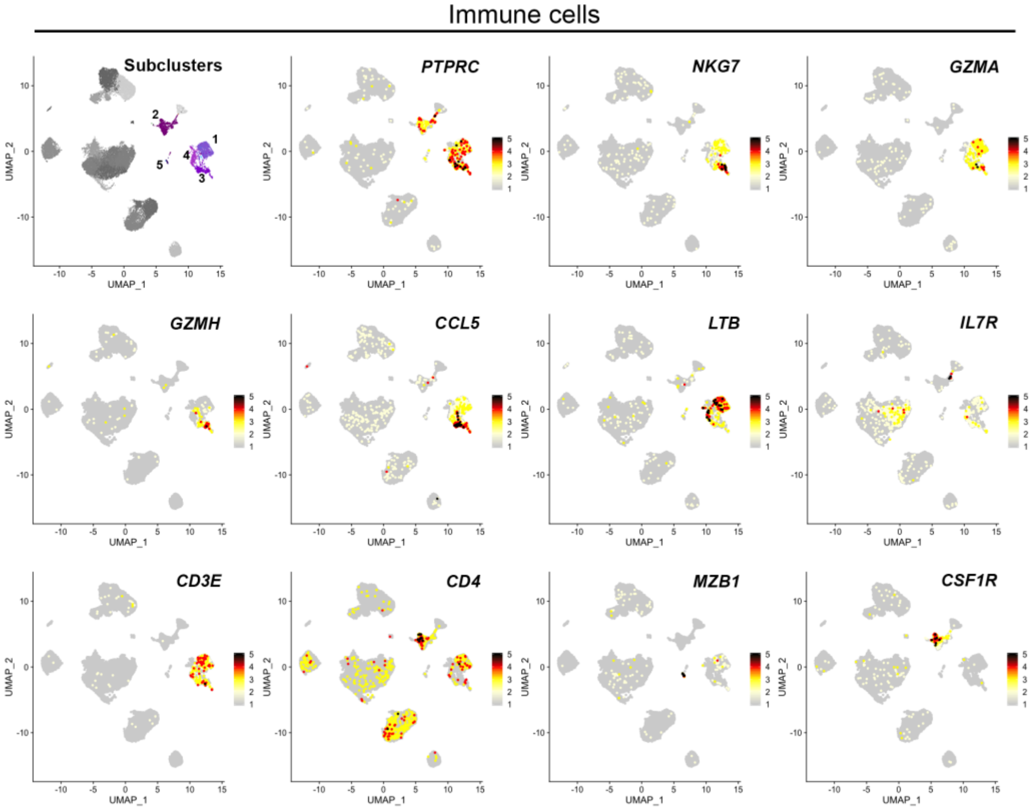


**Fig. S5. Feature plots showing the expression of genes characterizing specific subclusters of dental pulp immune cells.** All immune cells subclusters expressed the immune cells marker PTPRC (CD45). Clusters 1, 3 and 4 included T-cells and natural killer cells, as indicated by the expression of CD3E, CD4, GZMH, GZMA, and NKG67. Cluster 2 included macrophages and monocytes, as indicated by the high expression of CSF1R. Cluster 5 included B cells and plasma cells, as indicated by the expression of MZB1 and CD22 (see dataset) (Chetty and Gatter, 1994; Chitu and Stanley, 2006).

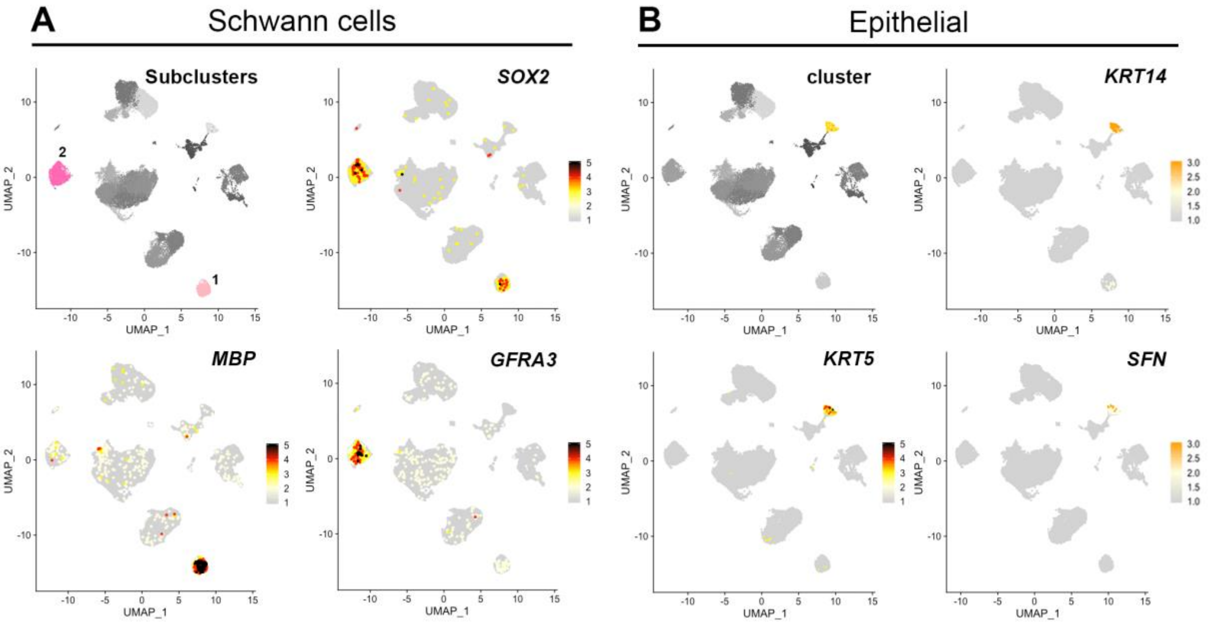


**Fig. S6. Feature plots showing the expression of genes characterizing subclusters of dental pulp Schwann cells (A) and epithelial cells (B).** A) All dental pulp Schwann cells express SOX2. MBP (myelin basic protein) is expressed by myelinating Schwann cells, while GFRA3 (GDNF family receptor alpha-3) marks non-myelinating Schwann cells. B) Epithelial cells express Keratin-coding genes, such as KRT14 and KRT5, as well as Stratifin (SFN). Keratins are intermediate filaments, and their expression is mostly restricted to epithelial cells (Herrmann et al., 2007; Karantza, 2011). SFN is expressed by differentiated keratinocytes, and it induces activation of adjacent fibroblasts by triggering expression of metalloproteases (Medina et al., 2007).

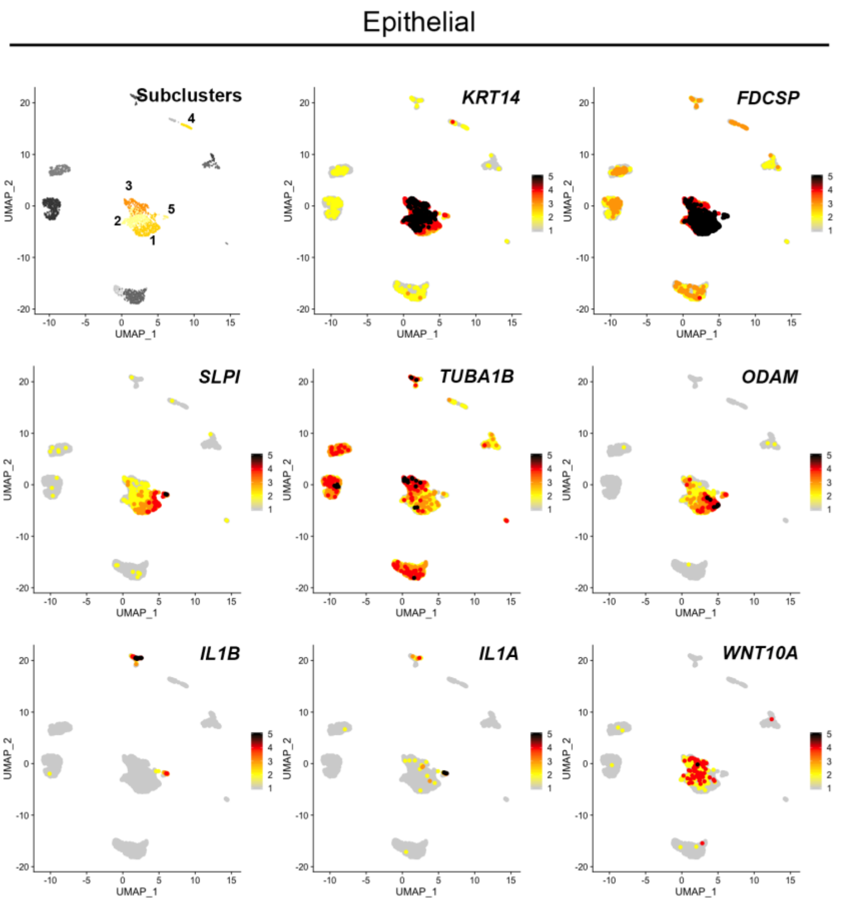


**Fig. S7. Feature plots showing the expression of genes characterizing subclusters of periodontal epithelial cells.** Epithelial cells represent the most abundant cell type in the human periodontium. Epithelial cells have been detected previously in mouse and human periodontal tissues (Athanassiou-Papaefthymiou et al., 2015; Tsunematsu et al., 2016). 5 subclusters could be identified. All epithelial cells express keratin-coding genes such as KRT14, and FDCSP (Follicular Dendritic Cells Secreted Protein). KRT14 is a common marker for dental epithelial cells (Tabata et al., 1996). FDCSP increases cell proliferation and inhibits the expression of genes associated with mineralization processes in periodontal MSCs of human teeth (Wei et al., 2011). Subcluster 1 was characterized by the expression of SLPI (secretory leukocyte protease inhibitor) and ODAM (Odontogenic Ameloblast-associated Protein). ODAM is expressed by periodontal epithelial cells that display stem cell properties (Athanassiou-Papaefthymiou et al., 2015). Clusters 2 and 3 showed higher expression of TUBA1B (tubulin alpha-1B chain) and WNT10A. WNT10A expression in dental epithelium is fundamental for tooth development and root formation (Mues et al., 2014; Yamashiro et al., 2007; Yu et al., 2020; Zhang et al., 2014). Cluster 5 showed higher expression of IL1A and IL1B, which are fundamental mediators of the immune response against infections (Miller and Cho, 2011), and they are strongly involved in the resolution of periodontal pathologies (Grigoriadou et al., 2010).

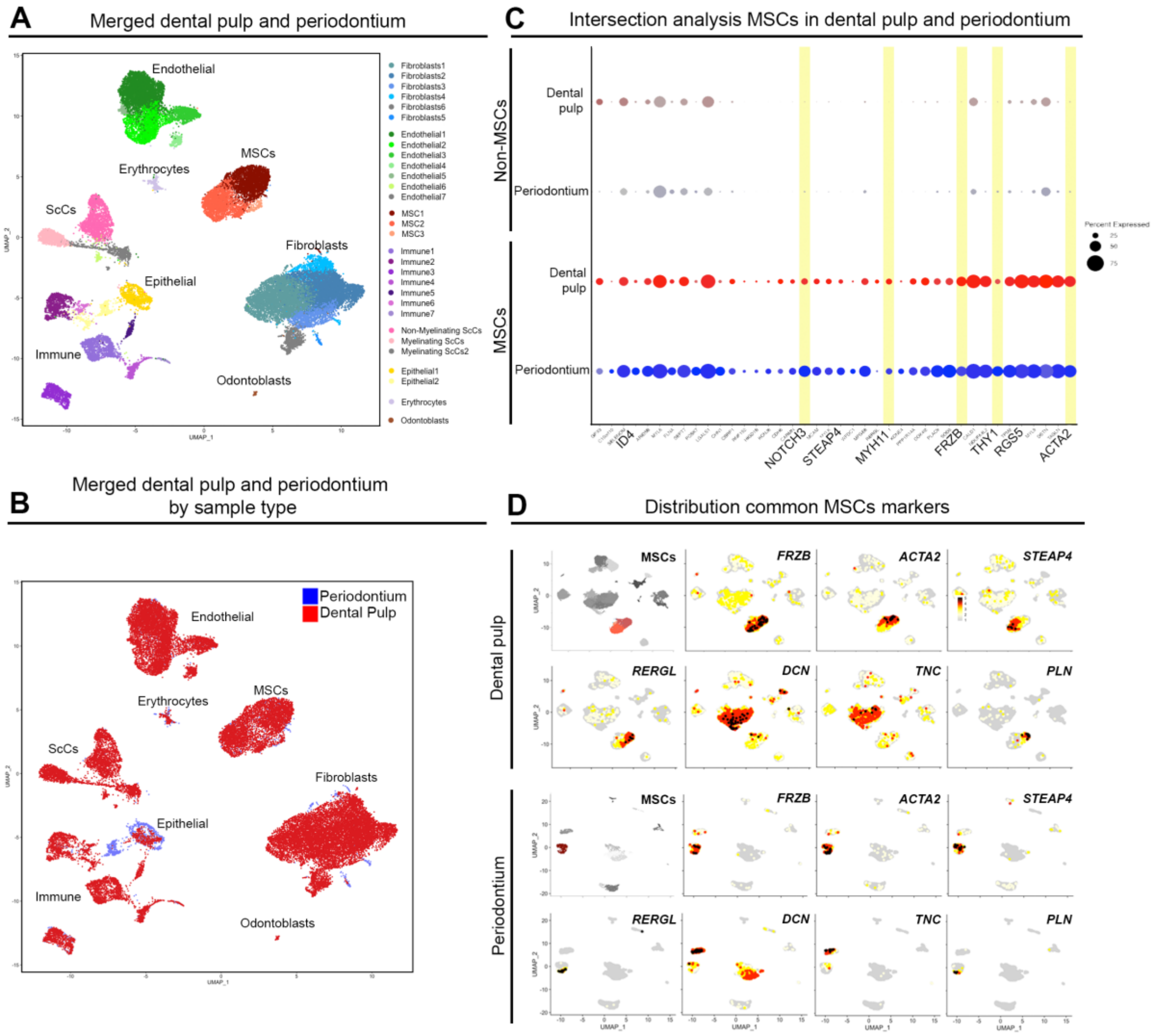


**Fig. S8. Comparison of dental pulp and periodontal MSCs.** A, B) UMAP plot showing the clusters distribution in the merged dental pulp / periodontium dataset. C) Dot plot showing the top 40 genes that characterize both dental pulp and periodontal MSCs against other dental cell types. Light yellow highlights genes of particular interest. MSCs in the dental pulp and the periodontium shared the expression of many stem cell markers and genes associated to stem cell function. MYH11 codes for a myosin heavy chain and its expression has been primarily observed in perivascular smooth muscle cells and pericytes, a common source of MSCs (Murgai et al., 2017). Similarly, ACTA2 is often expressed in pericytes. THY1 codes for CD90, a cell surface protein used as a classical marker for MSCs (An et al., 2018; Balic et al., 2010). MCAM/CD164 is a classical marker of MSCs in dental and non-dental tissues (Shi and Gronthos, 2003). RGS5 expression marks the perivascular NOTCH3+ MSCs in the dental pulp (Lovschall et al., 2007). Most MSCs populations express ID4, which codes for a transcription factor that inhibits cell differentiation (Junankar et al., 2015; Patel et al., 2015). D) Feature plots showing the distribution of the expression of common genes characterizing dental pulp and periodontal MSCs. FRZB is expressed by all MSCs, both in the dental pulp and in the periodontium. ACTA2, RERGL and PLN (Phospholamban) are particularly enriched in the MYH11+ MSCs subcluster, while DCN (Decorin) and STEAP4 are highly expressed in the THY1+ MSCs subcluster. TNC (Tenascin) is highly expressed in the CCL2+ MSCs subcluster. Previous studies have shown that TNC is expressed during odontogenesis in the dental mesenchyme (Vainio et al., 1989) as well as in the mature periodontium at the interface with cementum and with the alveolar bone (Lukinmaa et al., 1991; Midwood et al., 2016).

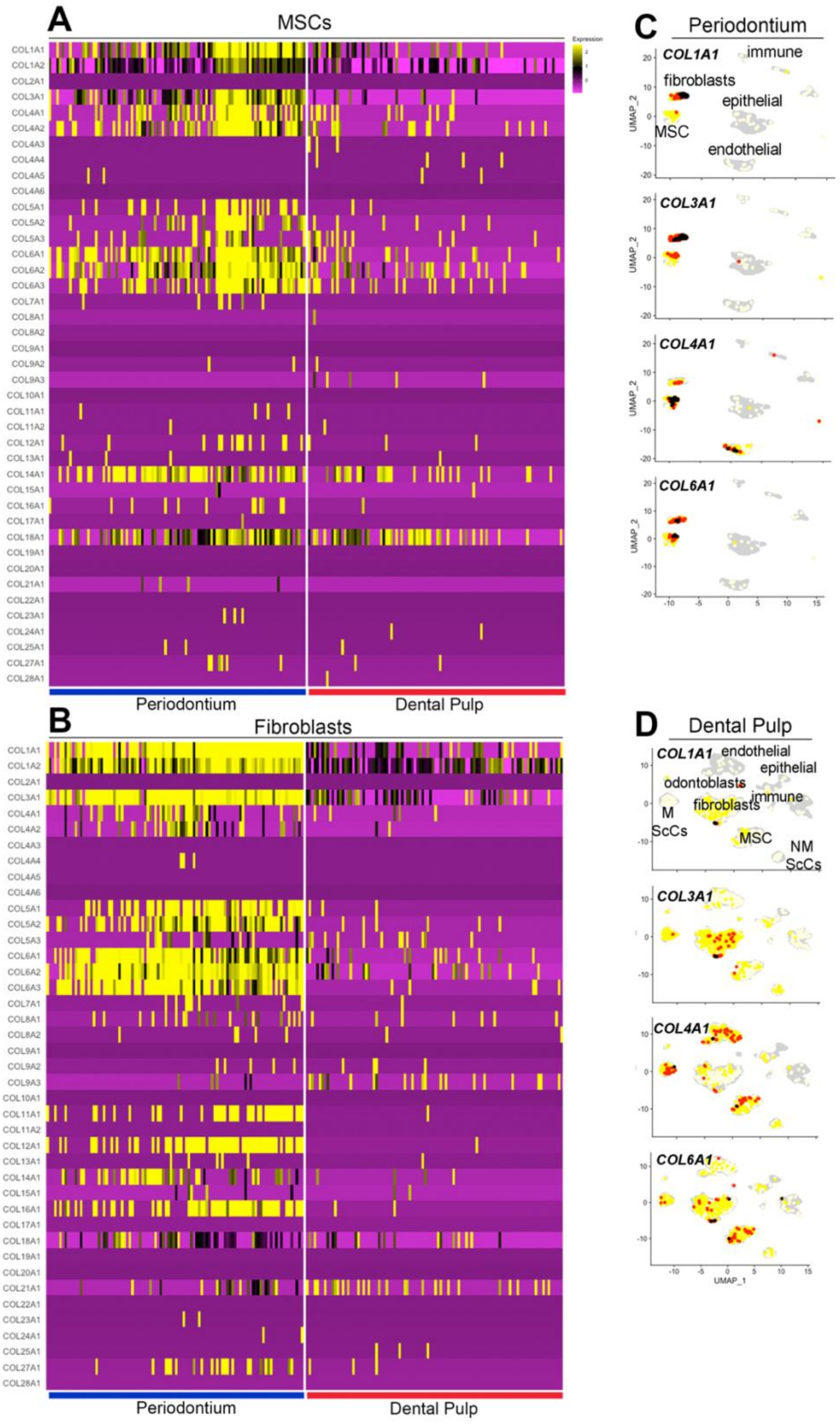


**Fig. S9. Expression of collagen-encoding genes.** A) Heatmap showing differential expression of genes encoding for collagens in MSCs from the periodontium and the dental pulp. B) Heatmap showing differential expression of genes encoding for collagens in fibroblasts from the periodontium and the dental pulp. C, D) Feature-plots showing the distribution of collagen-encoding genes in the periodontium (C) and in the dental pulp (D). Collagen-encoding genes are overall more expressed in the periodontium, in accordance to the intense remodeling that this tissue undergoes in response to mastication.

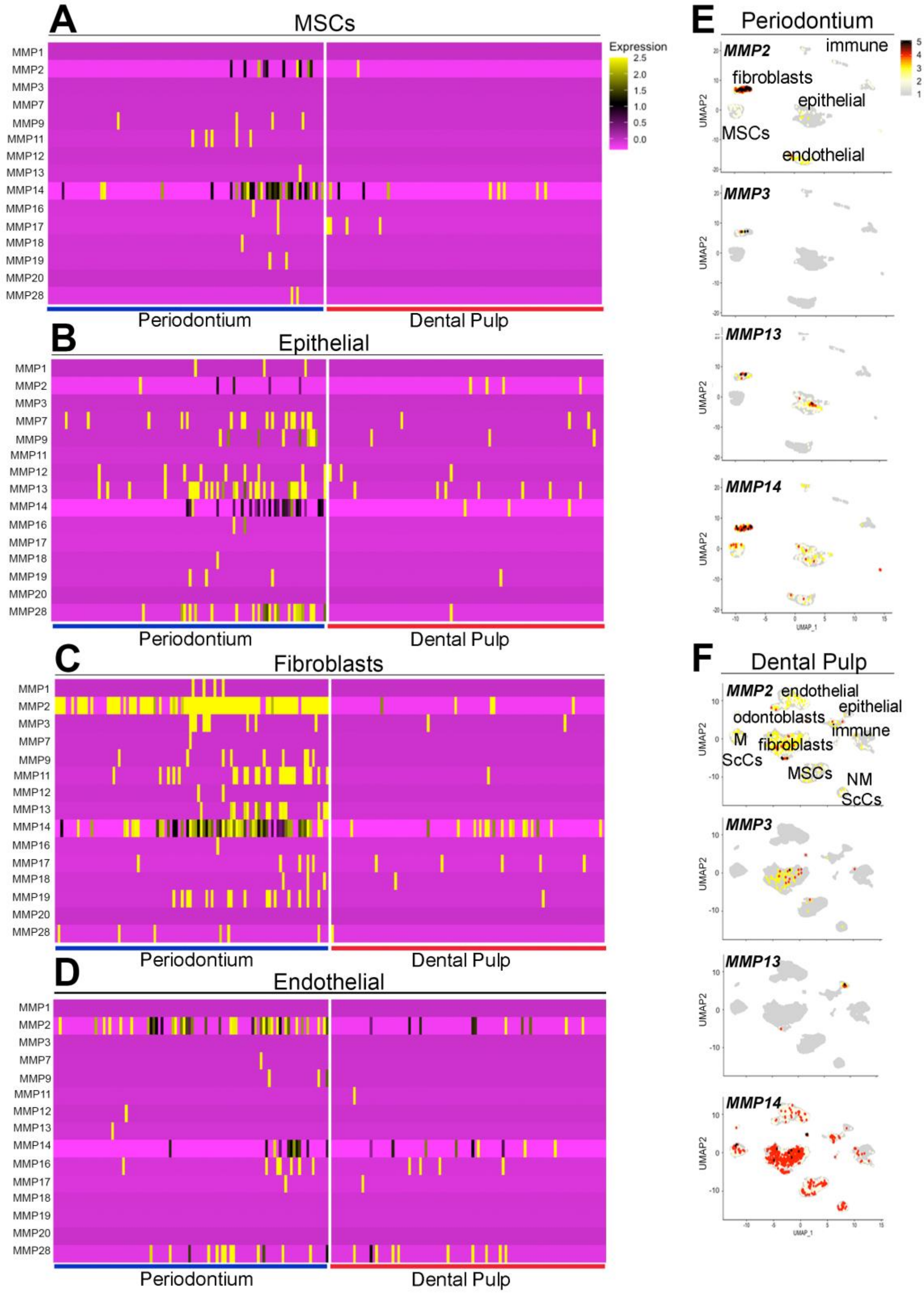


**Fig. S10. Expression of genes encoding metalloproteinases (MMPs).** A-D) Heatmap showing differential expression of genes encoding for MMPs in A) MSCs, B) epithelial cells, C) fibroblasts, D) endothelial cells, from the periodontium and the dental pulp. E, F) Feature-plots showing the distribution of collagen-encoding genes in the periodontium (E) and in the dental pulp (F). MMPs are actively involved in the turnover of the periodontal space since they degrade collagen and most of the secreted proteins that compose the periodontal extracellular matrix (Birkedal-Hansen, 1993; Sapna et al., 2014). MMPs-encoding genes are overall more expressed in the periodontium, in accordance with the intense remodeling of the extracellular matrix needed to compensate the stimuli that this tissue receives in response to mastication.

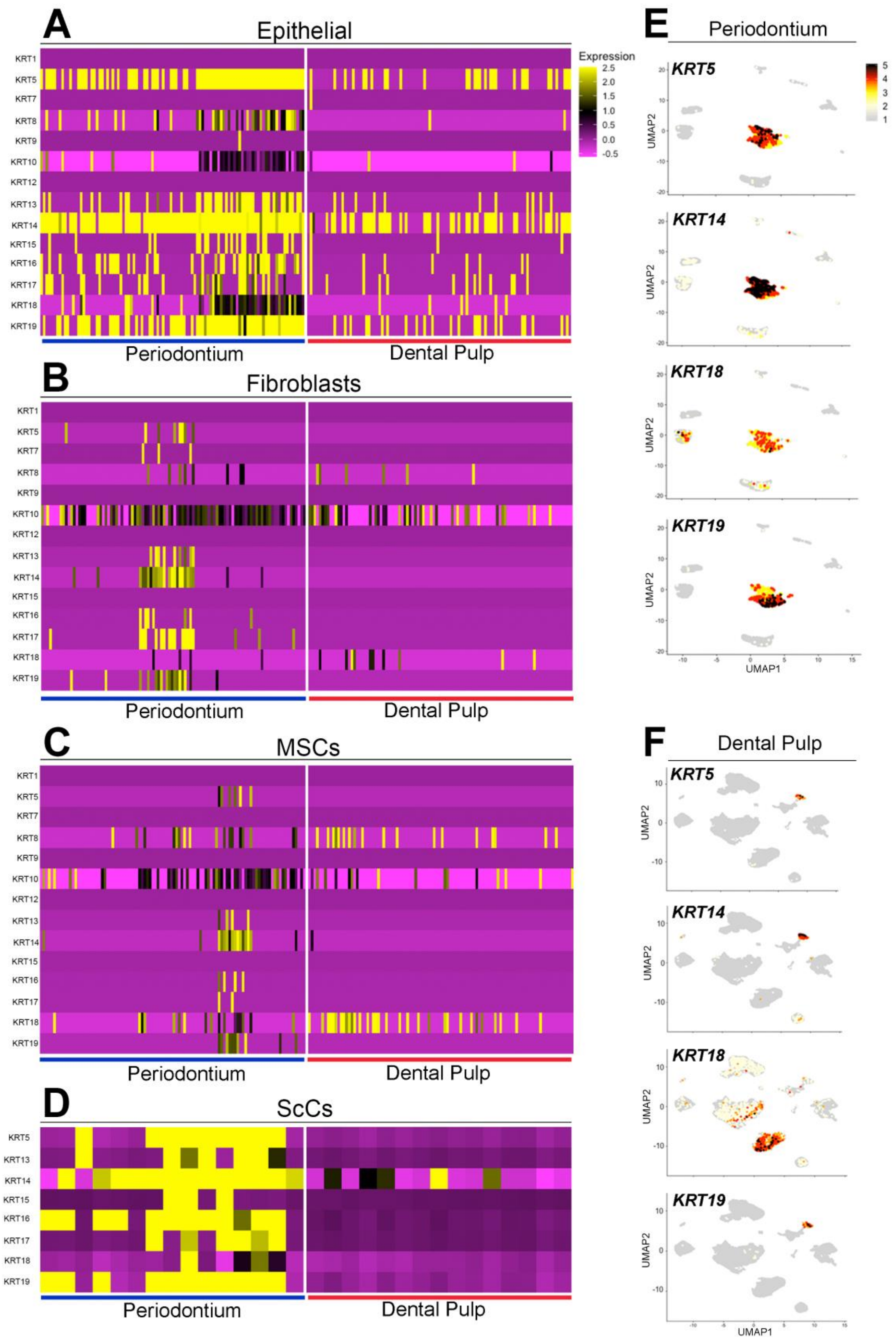


**Fig. S11. Expression of genes encoding keratins.** A-D) Heatmap showing differential expression of genes encoding for keratins in A) epithelial cells, B) fibroblasts, C) MSCs, D) Schwann cells from the periodontium and the dental pulp. E, F) Feature-plots showing the distribution of Keratin-encoding genes in the periodontium (E) and in the dental pulp (F). In the periodontium, Keratin-encoding genes are expressed also in non-epithelial cells. KRT18, a gene previously reported to be exclusively expressed in cells of single-layered and pseudostratified epithelia (Karantza, 2011), is expressed by MSCs in the dental pulp.

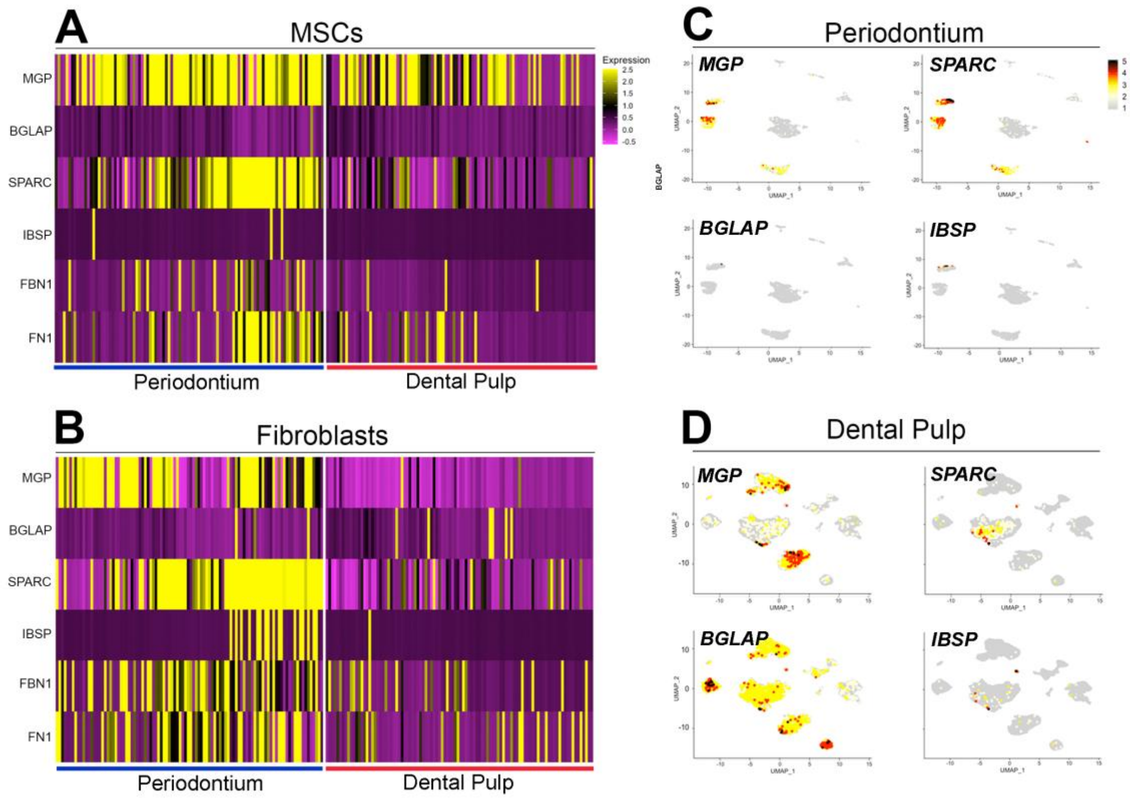


**Fig. S12. Expression of genes encoding non-collagenous bone-associated proteins.** A, B) Heatmaps showing genes differentially expressed between dental pulp and periodontal (A) MSCs and (B) fibroblasts. C, D) Feature plots showing the distribution of gene encoding non-collagenous bone-associated proteins in (C) periodontium and (D) dental pulp. Periodontal MSCs expressed higher levels of Osteonectin (SPARC) and MGP (Matrix Gla Protein) compared to dental pulp MSCs. Osteonectin is known to regulate Ca2+ deposition during bone formation (Termine et al., 1981), but in the periodontium its function is fundamental for proper collagen turnover and organization (Trombetta and Bradshaw, 2010). MGP (Matrix Gla Protein) is a potent inhibitor of mineralization (Kaipatur et al., 2008). Periodontal fibroblasts express higher levels of SPARC and MGP, as well as Osteocalcin (BGLAP) and Bone Sialophosphoprotein (BSP).

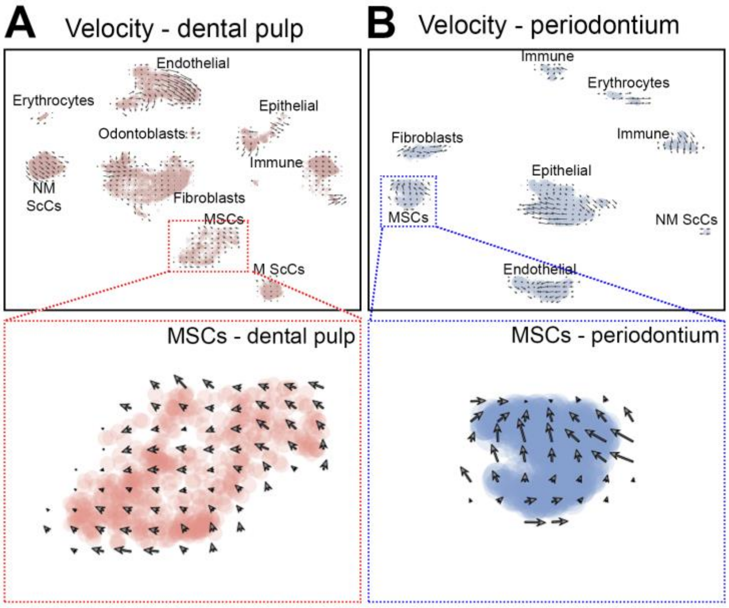


**Fig. S13. Velocity study of cell trajectories in the dental tissues.** A) Velocity in the dental pulp. B) Velocity in the periodontium. Red and blue rectangles highlight respectively the dental pulp and periodontal MSCs clusters shown in the global velocity plots. Only patients analyzed with 10X Genomics v3 kit were used in the velocity estimates.

Differential gene expression - Periodontium vs dental pulp MSCs

| | gene | p_val | avg_logFC | pct.1 | pct.2 | p_val_adj |
|---|---|---|---|---|---|---|
| 1 | HBB | 0 | 3.24581697177998 | 0.907 | 0.076 | 0 |
| 2 | HBA2 | 0 | 2.91169954034701 | 0.693 | 0.034 | 0 |
| 3 | HBA1 | 0 | 2.20113759332632 | 0.544 | 0.01 | 0 |
| 4 | NNMT | 0 | 1.87850058250694 | 0.629 | 0.02 | 0 |
| 5 | HOPX | 0 | 1.14458527446871 | 0.495 | 0.025 | 0 |
| 6 | RPL6P27 | 6.67256429342861E-267 | 1.11171714555118 | 0.742 | 0.106 | 1.65993381927623E-262 |
| 7 | FDCSP | 5.94640915612883E-262 | 2.75087526797022 | 0.307 | 0.006 | 1.47928820577017E-257 |
| 8 | PTMAP5 | 1.66832245588792E-243 | 1.14528932922911 | 0.633 | 0.079 | 4.15028577351238E-239 |
| 9 | COL3A1 | 7.52335353873889E-215 | 1.65707631127883 | 0.699 | 0.138 | 1.87158465983207E-210 |
| 10 | COL1A1 | 1.07439383971557E-213 | 1.59914236139218 | 0.751 | 0.162 | 2.67276955506042E-209 |
| 11 | SPARCL1 | 2.99317807053905E-189 | 1.29053911941933 | 0.909 | 0.287 | 7.44612908607998E-185 |
| 12 | CXCL12 | 4.2262140200075E-156 | 1.19411865882346 | 0.443 | 0.064 | 1.05135526175727E-151 |
| 13 | IGFBP4 | 3.92974692001991E-147 | 1.02031640578958 | 0.68 | 0.171 | 9.77603141293352E-143 |
| 14 | S100A4 | 4.38097554588702E-137 | 1.08885329373367 | 0.915 | 0.393 | 1.08985528655031E-132 |
| 15 | DCN | 3.21450625730978E-126 | 1.17058089254281 | 0.67 | 0.198 | 7.99672721630954E-122 |
| 16 | COL6A1 | 1.9531483174306E-124 | 1.02344023915478 | 0.68 | 0.191 | 4.8588470692721E-120 |
| 17 | APOE | 2.62528933909972E-108 | 1.48546734763149 | 0.695 | 0.245 | 6.53093228887836E-104 |
| 18 | CXCL14 | 2.21414963747453E-89 | -2.04411843929188 | 0.151 | 0.606 | 5.50814005314539E-85 |
| 19 | SPARC | 1.04752798714087E-85 | 1.00238177918575 | 0.862 | 0.481 | 2.60593537361035E-81 |
| 20 | IFI27 | 6.94621489604408E-69 | -1.58810329223689 | 0.328 | 0.635 | 1.72800987968888E-64 |
| 21 | TF | 1.08038343540846E-45 | -1.51444432023821 | 0.012 | 0.325 | 2.68766987226564E-41 |
| 22 | KRT18 | 1.57589643449871E-39 | -1.46872929126273 | 0.157 | 0.428 | 3.92035756010244E-35 |
| 23 | PTN | 2.35938566952991E-39 | -1.32588047483401 | 0.414 | 0.588 | 5.86944373008955E-35 |
| 24 | CLU | 5.03668354194873E-29 | -1.06405551400471 | 0.212 | 0.424 | 1.25297576473059E-24 |
| 25 | IFI6 | 3.16052123818713E-28 | -1.13704670755114 | 0.334 | 0.497 | 7.86242868423814E-24 |
| 26 | ISG15 | 5.16318354573924E-27 | -1.28475423457449 | 0.219 | 0.415 | 1.28444517067355E-22 |
| 27 | RBP1 | 1.78272016950256E-24 | -1.05038498240697 | 0.058 | 0.253 | 4.43487296567152E-20 |
| 28 | RARRES1 | 2.5228534697408E-21 | -1.00182017404775 | 0.008 | 0.172 | 6.27610257667419E-17 |
| 29 | PTGDS | 5.07390837482826E-20 | -1.01012662211226 | 0.027 | 0.189 | 1.26223618640603E-15 |
| 30 | IGFBP6 | 6.09168123937234E-19 | -1.0714522518631 | 0.264 | 0.403 | 1.51542754191866E-14 |
| 31 | RHOB | 6.15620502917967E-15 | -1.03599434939694 | 0.328 | 0.419 | 1.53147912510903E-10 |
| 32 | PPP1CB | 7.01853477480945E-11 | -1.06066566583755 | 0.454 | 0.465 | 1.74600089592935E-06 |
| 33 | DDIT4 | 1.0355171449118E-10 | -1.03502796781716 | 0.198 | 0.302 | 2.5760560013971E-06 |

**Table S1. Genes differentially expressed (Fc > 1; p < 0.005) between periodontal and dental pulp MSCs.** Periodontal MSCs expressed higher levels of CCL2 and Collagen encoding genes. Periodontal MSCs were also characterized by higher expression of SPARC/Osteonectin, a secreted molecule fundamental for the regulation of periodontal homeostasis and Collagen content (Trombetta and Bradshaw, 2010). Dental pulp MSCs expressed higher levels of CXCL14, is associated with increased angiogenic potential (Hayashi et al., 2015), and RARRES1 which mediates retinoic acid-responses in stem cells (Oldridge et al., 2013). Dental pulp MSCs strongly expressed KRT18, a gene previously reported to be exclusively expressed in cells of single-layered and pseudostratified epithelia (Omary et al., 2009).

### Pairwise extended jaccard similarity - ranks

| | Cell type 1 | Cell type 2 |
|---|---|---|
| 1 | Epithelial_perio | Myelinating_ScCs perio |
| 2 | Endothelial_perio | Endothelial_pulp |
| 3 | Erythrocytes_perio | Erythrocytes_pulp |
| 4 | MSC_perio | MSC_pulp |
| 5 | Non-Myelinating_ScCs_pulp | Myelinating_ ScCs_pulp |
| 6 | Epithelial_pulp | Myelinating_ScCs_perio |
| 7 | Odontoblasts_pulp | Fibroblasts_pulp |
| 8 | Fibroblasts_perio | Cementoblasts_perio |
| 9 | Non-Myelinating ScCs_perio | Non-Myelinating_ ScCs_pulp |
| 10 | Immune_perio | Immune_pulp |

### Number of DEG - cell type

| | cell types | Number of DEG |
|---|---|---|
| 1 | MSC | 333 |
| 2 | Fibroblasts | 688 |
| 3 | Erythrocytes | 125 |
| 4 | Epithelial | 545 |
| 5 | Immune | 351 |
| 6 | Endothelial | 475 |
| 7 | Non-Myelinating ScCs | 143 |
| 8 | Odontoblasts / Cementoblasts | 107 |
| 9 | Myelinating ScCs | 313 |

**Table S2.** Left: Periodontal and dental pulp cell types ranked according to pairwise extended jaccard similarity. Right: number of differentially expressed genes between equivalent cell types in the dental pulp and the periodontium

### Collagen-encoding genes

(Periodontium vs dental pulp)

#### Fibroblasts

| | gene | p_val | avg_logFC | pct.1 | pct.2 | p_val_adj |
|---|---|---|---|---|---|---|
| 111 | COL1A1 | 2.25487242834546E-166 | 3.3019368581082 | 0.928 | 0.376 | 5.60944613999501E-162 |
| 189 | COL1A2 | 5.24386883102724E-106 | 1.96956844269406 | 0.975 | 0.737 | 1.30451724909465E-101 |
| 120 | COL3A1 | 2.22198351415981E-153 | 2.40575302329749 | 0.957 | 0.495 | 5.52762838817536E-149 |
| 105 | COL4A1 | 7.45156869789291E-180 | 0.83048292325594 | 0.343 | 0.027 | 1.85372674497482E-175 |
| 123 | COL4A2 | 1.71151093270727E-150 | 0.712807129120808 | 0.379 | 0.04 | 4.25772574729587E-146 |
| 17 | COL5A1 | 0 | 1.22445080957512 | 0.632 | 0.03 | 0 |
| 75 | COL5A2 | 3.05331225252665E-254 | 1.23286104790791 | 0.733 | 0.092 | 7.59572489061056E-250 |
| 72 | COL6A1 | 9.39035086594511E-259 | 1.74716413138289 | 0.866 | 0.144 | 2.33603758492117E-254 |
| 89 | COL6A2 | 3.43727039099989E-206 | 1.89125151813711 | 0.924 | 0.241 | 8.55089755169042E-202 |
| 6 | COL6A3 | 0 | 1.82762970827482 | 0.834 | 0.075 | 0 |
| 635 | COL9A3 | 1.03141268744368E-09 | -0.643726096050555 | 0.043 | 0.18 | 2.56584534255364E-05 |
| 16 | COL11A1 | 0 | 1.27768783976508 | 0.444 | 0.013 | 0 |
| 12 | COL12A1 | 0 | 1.53579904694227 | 0.61 | 0.006 | 0 |
| 20 | COL14A1 | 0 | 1.1029391222703 | 0.455 | 0.019 | 0 |
| 23 | COL16A1 | 0 | 1.02744676887705 | 0.538 | 0.026 | 0 |
| 476 | COL18A1 | 2.47330333647109E-32 | 0.254176480519037 | 0.419 | 0.139 | 6.15283671013913E-28 |
| 937 | COL21A1 | 0.115313439128048 | -0.484816613063316 | 0.213 | 0.219 | 1 |

#### MSCs

| | gene | p_val | avg_logFC | pct.1 | pct.2 | p_val_adj |
|---|---|---|---|---|---|---|
| 12 | COL1A1 | 1.07439383971557E-213 | 1.59914236139218 | 0.751 | 0.162 | 2.67276955506042E-209 |
| 189 | COL1A2 | 4.7454973002183E-46 | 0.728211899450018 | 0.732 | 0.399 | 1.18053736337531E-41 |
| 11 | COL3A1 | 7.52335353873889E-215 | 1.65707631127883 | 0.699 | 0.138 | 1.87158465983207E-210 |
| 53 | COL4A1 | 8.8143610293432E-103 | 0.860257329496067 | 0.495 | 0.118 | 2.19274859326971E-98 |
| 120 | COL4A2 | 3.05435194531821E-66 | 0.568640888255137 | 0.555 | 0.192 | 7.59831133436812E-62 |
| 60 | COL5A1 | 6.68949374408408E-95 | 0.327013912144249 | 0.216 | 0.021 | 1.6641453587158E-90 |
| 92 | COL5A2 | 2.17400695967927E-76 | 0.484278088748829 | 0.338 | 0.07 | 5.40827711359411E-72 |
| 32 | COL6A1 | 1.9531483174306E-124 | 1.02344023915478 | 0.68 | 0.191 | 4.8588470692721E-120 |
| 72 | COL6A2 | 4.42759676271991E-89 | 0.831620893206911 | 0.744 | 0.292 | 1.10145324666183E-84 |
| 41 | COL6A3 | 2.15934421027835E-115 | 0.706814172812724 | 0.532 | 0.119 | 5.37180059190944E-111 |
| 105 | COL14A1 | 9.36381906188564E-70 | 0.505434369436353 | 0.647 | 0.227 | 2.32943726802529E-65 |
| 57 | COL16A1 | 6.58947716426885E-100 | 0.344835214148013 | 0.169 | 0.01 | 1.63926423415516E-95 |

**Table S3.** Collagen-encoding genes differentially expressed ($p < 0.001$) between periodontal and dental pulp fibroblasts and MSCs.

### Keratin-encoding genes

(Periodontium vs dental pulp)

#### Epithelial

| | gene | p_val | avg_logFC | pct.1 | pct.2 | p_val_adj |
|---|---|---|---|---|---|---|
| 51 | KRT5 | 1.7760833374321E-50 | 0.662134563892527 | 0.773 | 0.269 | 4.41836251852985E-46 |
| 21 | KRT8 | 1.95430325171363E-55 | 0.296545123782922 | 0.423 | 0.035 | 4.861720199288E-51 |
| 117 | KRT13 | 2.40526568560614E-37 | 0.489124101031112 | 0.478 | 0.115 | 5.9835794460824E-33 |
| 104 | KRT14 | 9.53489912531268E-40 | 0.512019219754885 | 0.954 | 0.551 | 2.37199685540404E-35 |
| 237 | KRT15 | 1.06289366570786E-21 | 0.327423789729451 | 0.211 | 0.03 | 2.64416057218144E-17 |
| 27 | KRT18 | 2.37494190777221E-54 | 0.371834626953621 | 0.466 | 0.057 | 5.90814298396493E-50 |
| 15 | KRT19 | 2.91730913531038E-57 | 0.770694899057093 | 0.792 | 0.272 | 7.25738993591163E-53 |
| 211 | KRT8 | 1.95430325171363E-55 | 0.296545123782922 | 0.423 | 0.035 | 4.861720199288E-51 |

#### MSCs

| | gene | p_val | avg_logFC | pct.1 | pct.2 | p_val_adj |
|---|---|---|---|---|---|---|
| 306 | KRT8 | 8.17811740716836E-12 | -0.788924436402796 | 0.107 | 0.224 | 2.03447026738127E-07 |
| 56 | KRT14 | 1.02735136523106E-100 | 0.489689085742613 | 0.132 | 0.004 | 2.55574199128531E-96 |
| 220 | KRT18 | 1.57589643449867E-39 | -1.46872929126273 | 0.157 | 0.428 | 3.92035756010235E-35 |
| 3061 | KRT8 | 8.17811740716836E-12 | -0.788924436402796 | 0.107 | 0.224 | 2.03447026738127E-07 |

#### Fibroblasts

| | gene | p_val | avg_logFC | pct.1 | pct.2 | p_val_adj |
|---|---|---|---|---|---|---|
| 33 | KRT14 | 0 | 0.774716256198696 | 0.224 | 0.003 | 0 |
| 63 | KRT17 | 2.6869875919885E-296 | 0.310658566332385 | 0.148 | 0.001 | 6.6844190325898E-292 |

#### ScCs

| | gene | p_val | avg_logFC | pct.1 | pct.2 | p_val_adj |
|---|---|---|---|---|---|---|
| 10 | KRT5 | 9.49024512712327E-177 | 2.30252721192442 | 0.538 | 0.002 | 2.36088828027446E-172 |
| 125 | KRT8 | 3.98619631248065E-32 | 0.803345311854587 | 0.333 | 0.02 | 9.91646056655812E-28 |
| 903 | KRT10 | 0.136263579018209 | -0.275482038836599 | 0.41 | 0.25 | 1 |
| 22 | KRT13 | 4.97856780968926E-141 | 2.10478124975943 | 0.359 | 0 | 1.2385183140164E-136 |
| 156 | KRT14 | 1.58832510524415E-20 | 2.80853628699739 | 0.667 | 0.177 | 3.95127636431588E-16 |
| 95 | KRT15 | 2.5345804340202E-51 | 0.326997146876151 | 0.128 | 0 | 6.30527574571206E-47 |
| 16 | KRT16 | 2.05969189935912E-161 | 2.31989135139677 | 0.436 | 0.001 | 5.12389553803569E-157 |
| 75 | KRT17 | 1.72889343997505E-62 | 0.77870053053477 | 0.231 | 0.002 | 4.30096821062594E-58 |
| 193 | KRT18 | 2.92626668227313E-13 | 0.498738700096318 | 0.282 | 0.037 | 7.27967362549086E-09 |
| 5 | KRT19 | 7.38677820170057E-193 | 2.87943743216864 | 0.538 | 0.001 | 1.83760881323705E-188 |
| 1251 | KRT8 | 3.98619631248065E-32 | 0.803345311854587 | 0.333 | 0.02 | 9.91646056655812E-28 |

**Table S4.** Keratin-encoding genes differentially expressed ($p < 0.001$) between periodontal and dental pulp epithelial cells, MSCs, fibroblasts and ScCs.

**MMP-encoding genes**

(Periodontium vs dental pulp)

Fibroblasts

| | gene | p_val | avg_logFC | pct.1 | pct.2 | p_val_adj |
|---|---|---|---|---|---|---|
| 67 | MMP2 | 4.6148452189556E-282 | 1.9538842190864 | 0.83 | 0.124 | 1.14803504511958E-277 |
| 118 | MMP11 | 3.64223980263109E-156 | 0.265536622985388 | 0.206 | 0.01 | 9.06079995700536E-152 |
| 11 | MMP13 | 0 | 1.54239457203761 | 0.191 | 0 | 0 |
| 319 | MMP14 | 5.78794888219009E-62 | 0.471100307125854 | 0.61 | 0.178 | 1.43986804342243E-57 |

Epithelial

| | gene | p_val | avg_logFC | pct.1 | pct.2 | p_val_adj |
|---|---|---|---|---|---|---|
| 256 | MMP7 | 1.93984660931978E-20 | 0.335364772334984 | 0.196 | 0.027 | 4.82575641000482E-16 |
| 423 | MMP12 | 2.90533031306277E-11 | 0.319730178933507 | 0.129 | 0.027 | 7.22759021980626E-07 |
| 175 | MMP13 | 1.55228121633805E-26 | 0.553801448188468 | 0.28 | 0.049 | 3.86160998188416E-22 |

Endothelial

| | gene | p_val | avg_logFC | pct.1 | pct.2 | p_val_adj |
|---|---|---|---|---|---|---|
| 7 | MMP2 | 1.30077357343548E-131 | 0.595358740089258 | 0.386 | 0.066 | 3.23593441863545E-127 |

**Table S5.** Genes encoding for metalloproteases (MMP) differentially expressed ($p < 0.001$) between periodontal and dental pulp fibroblasts and MSCs.

### Non collagenous bone-associated proteins
(periodontium vs dental pulp)

#### Fibroblasts

| | gene | p_val | avg_logFC | pct.1 | pct.2 | p_val_adj |
|---|---|---|---|---|---|---|
| 81 | MGP | 1.78588906938445E-219 | 1.99197028249637 | 0.805 | 0.141 | 4.4427562379077E-215 |
| 213 | SPARC | 5.16602480192375E-94 | 1.94575060495438 | 0.877 | 0.515 | 1.28515198997457E-89 |
| 73 | IBSP | 1.93163964632213E-255 | 1.46661576267429 | 0.181 | 0.003 | 4.80533994815555E-251 |
| 343 | FBN1 | 7.1117677680317E-58 | 0.533218478334099 | 0.625 | 0.192 | 1.76919446765325E-53 |
| 330 | FN1 | 2.98712269762832E-60 | 0.581968071589218 | 0.632 | 0.194 | 7.43106513488997E-56 |

#### MSCs

| | gene | p_val | avg_logFC | pct.1 | pct.2 | p_val_adj |
|---|---|---|---|---|---|---|
| 233 | MGP | 8.87543348342118E-37 | 0.501073027684103 | 0.874 | 0.545 | 2.20794158767069E-32 |
| 80 | SPARC | 1.04752798714087E-85 | 1.00238177918576 | 0.862 | 0.481 | 2.60593537361035E-81 |

**Table S6.** Genes encoding for non-collagenous bone associated-proteins differentially expressed ($p < 0.001$) between periodontal and dental pulp fibroblasts and MSCs.
